# The proximity-based protein interaction landscape of the transcription factor p65 NF-κB / RELA and its gene-regulatory logics

**DOI:** 10.1101/2024.01.03.574021

**Authors:** Lisa Leib, Jana Juli, Liane Jurida, Johanna Meier-Soelch, Christin Mayr-Buro, M. Lienhard Schmitz, Daniel Heylmann, Axel Weber, Argyris Papantonis, Marek Bartkuhn, Jochen Wilhelm, Uwe Linne, Michael Kracht

## Abstract

The protein interactome of p65 / RELA, the most active subunit of the transcription factor (TF) NF-κB, has not been previously determined in living cells. Using p65-miniTurbo fusion proteins, we identified by biotin tagging > 350 RELA interactors from untreated and IL-1α-stimulated cells, including many TFs (47 % of all interactors) and > 50 epigenetic regulators belonging to different classes of chromatin remodeling complexes. According to point mutants of p65, the interactions primarily require intact dimerization rather than DNA binding properties. A targeted RNAi screen for 38 interactors and subsequent functional transcriptome and bioinformatics studies identified gene regulatory (sub)networks, each controlled by RELA in combination with one of the TFs ZBTB5, GLIS2, TFE3 / TFEB or S100A8 / A9. The remarkably large, dynamic and versatile high resolution interactome of RELA and its gene-regulatory logics provides a rich resource and a new framework for explaining how RELA cooperativity determines gene expression patterns.

**Highlights:**

- Identification of > 350 largely dimerization-dependent interactors of p65 / RELA by miniTurboID
- The interactome is dominated by transcription factors and epigenetic regulator complexes
- Functional validation of 38 high confidence interactors by targeted siRNA screen
- Identification of genetic networks regulated by RELA and six of its interactors in the IL-1α response

## Introduction

Transcription factors (TFs) comprise a large family of proteins that read and interpret the genome to decode the DNA sequence. TFs are defined by their sequence–specific DNA binding domains (DBD) and by the ability to induce or repress transcription (Fulton et al., 2009; Vaquerizas et al., 2009). A recent combinatorial survey catalogued a total of 1639 human TFs (http://humantfs.ccbr.utoronto.ca) (Lambert et al., 2018).

TFs are key components of gene regulatory networks in which the spatio-temporal expression patterns of TFs and their auto-regulatory loops determine cell identity, developmental processes, and disease states (Almeida et al., 2021; Fuxman Bass et al., 2015).

So far, for only a limited number of TFs the (direct) binding to chromatin has been mapped comprehensively using ChIPseq or related techniques. However, bioinformatics analyses of sequence motifs deduced from genome-wide DNA footprints in open chromatin regions suggest that transcription factors generally bind cooperatively to enhancers or promoters to execute their gene regulatory functions (Funk et al., 2020; Neph et al., 2012; Vierstra et al., 2020).

In contrast to the wealth of information on (predicted) DNA binding of TFs, we currently lack a global understanding of TF protein–protein interactions (PPIs) and their functional contributions to TF cooperativity within transcriptional networks. A recent, large-scale study examined the basal interactomes of 109 common TFs to find, depending on the method applied, 1538 to 6703 PPI, respectively. This new evidence suggests that TF cooperativity may be largely determined through the repertoire of (dynamic) PPI (Goos et al., 2022).

The REL DBD is found in only 10 (0.6%) of all human TFs, including the five members of the nuclear factor of kappa light polypeptide gene enhancer in B-cells (NF-κB) family of TFs (Lambert et al., 2018). The five NF-kB subunits are evolutionary conserved, inducible regulators of development and disease conditions and are particularly important in regulating innate and adaptive immune responses (Williams and Gilmore, 2020; Zhang et al., 2017).

The p65 NF-κB/ RELA transcription factor contains a highly structured and conserved N-terminal REL homology domain (RHD) that is involved in DNA binding and dimerization, as shown by the crystal structure of the p65 / p50 NF-κBheterodimer bound to DNA (Chen et al., 1998; Williams and Gilmore, 2020). The C-terminal half of p65 / RELA contains two potent transactivation domains (TA_1_, TA_2_) and is inducibly phosphorylated at multiple residues (Christian et al., 2016; Viatour et al., 2005). In contrast to the RHD, the C-terminus is highly unstructured as revealed originally by NMR and supported by bioinformatics predictions (Jumper et al., 2021; Schmitz et al., 1994; Schmitz et al., 1995; Varadi et al., 2022). This phenomenon may also underly the previously reported cytokine–dependent conformational switches of p65 / RELA (Milanovic et al., 2014).

Both, post-translational modifications and structural flexibility of the p65 / RELA C-terminus are features that have very likely evolved to expand the repertoire of PPI under changing conditions and within subcellular compartments. However, the p65 / RELA interactome has not yet been determined comprehensively with methods that also cover labile, transient or sub-stoichiometric interactions of p65 / RELA with cellular proteins as they occur in living cells.

Accordingly, we have limited understanding of how exactly p65 / RELA cooperates with partner TFs (*in cis* at overlapping DNA elements or *in trans* across chromosomes), chromatin modifiers, the transcription machinery, and other nuclear cofactors to regulate transcription of specific groups of genes (Bacher et al., 2021).

Biotin-proximity labeling is a relatively new approach by which cells, expressing a bait protein fused to an engineered bacterial biotin ligase (such as BirA* used for BioID), are incubated with biotin for several hours to biotinylate all proteins in close vicinity (i.e. a radius of 1-10 nm) to the fusion protein. After cell lysis, biotinylated proteins are captured on streptavidin affinity matrices and subsequently identified by liquid chromatography-tandem mass spectrometry (LC-MS / MS) (Qin et al., 2021; Roux et al., 2012). Key advantages of this approach include its ability to capture weak or transient interactions from both soluble and insoluble proteins, or subcellular organelles, and the possibility to use high-stringency protein purification methods to reduce background contaminants (Zhou and Zou, 2021). MiniTurbo (mTb, used for miniTurboID) is a recently improved small 28 kDa variant of BirA that more efficiently biotinylates intracellular proteins within minutes in the presence of exogenously added biotin (Branon et al., 2018).

The pro-inflammatory cytokine interleukin-1 (IL-1) rapidly activates p65 / RELA in a broad range of cell types (Meier-Soelch et al., 2021). Numerous studies, including our own work, have shown the importance of p65 / RELA for the expression of IL-1-target genes, rendering this system an ideal model to study the dynamic p65 / RELA interactome (Barter et al., 2021; Jurida et al., 2015; Weiterer et al., 2020).

Here, we report the first proximity-based p65 / RELA interactome using inducible wild type, or mutant p65-miniTurbo fusion proteins devoid of either DNA binding or dimerization properties, to identify 366 high confidence p65 / RELA interactors (HCI) of which 87 % are novel. The p65 / RELA interactome is highly enriched for nuclear proteins, including 172 TFs (47 % of all p65 / RELA interactors) and 74 epigenetic regulators (20% of all interactors). A targeted siRNA screen for 38 interactors and subsequent functional gene expression studies identify new gene regulatory (sub)networks, each activated or repressed by p65 / RELA in combination with one of the TFs ZBTB5, GLIS2, TFE3 / TFEB or S100A8/ S100A9.

Taken together, these data reveal a remarkably large, dynamic and versatile (transcription factor and epigenetic regulator) interactome of p65, which determines gene expression patterns mainly via DNA-independent protein-protein interactions (PPIs).

## Results

### Identification of new p65 / RELA interactors using miniTurbo-based proximity labeling

Point mutations in the RHD of p65 / RELA, such as glutamate 39 to isoleucine (E / I), inhibit DNA binding, while mutations of two other amino acids, phenylalanine 213 and leucine 215, to aspartic acid (FL / DD), prevent dimerization as previously shown by co-immunoprecipitation and EMSA experiments (Riedlinger et al., 2019) **(Fig.1A, left image)**. In contrast, the C-terminal half of p65 / RELA is highly unstructured as revealed by alphafold (Jumper et al., 2021; Varadi et al., 2022) **(Fig.1A, right image)**. We reasoned that these features of the RHD and the p65 / RELA C-terminus very likely evolved to expand the repertoire of possible protein interactions of p65 / RELA under changing conditions and within subcellular compartments.

**Fig. 1.**
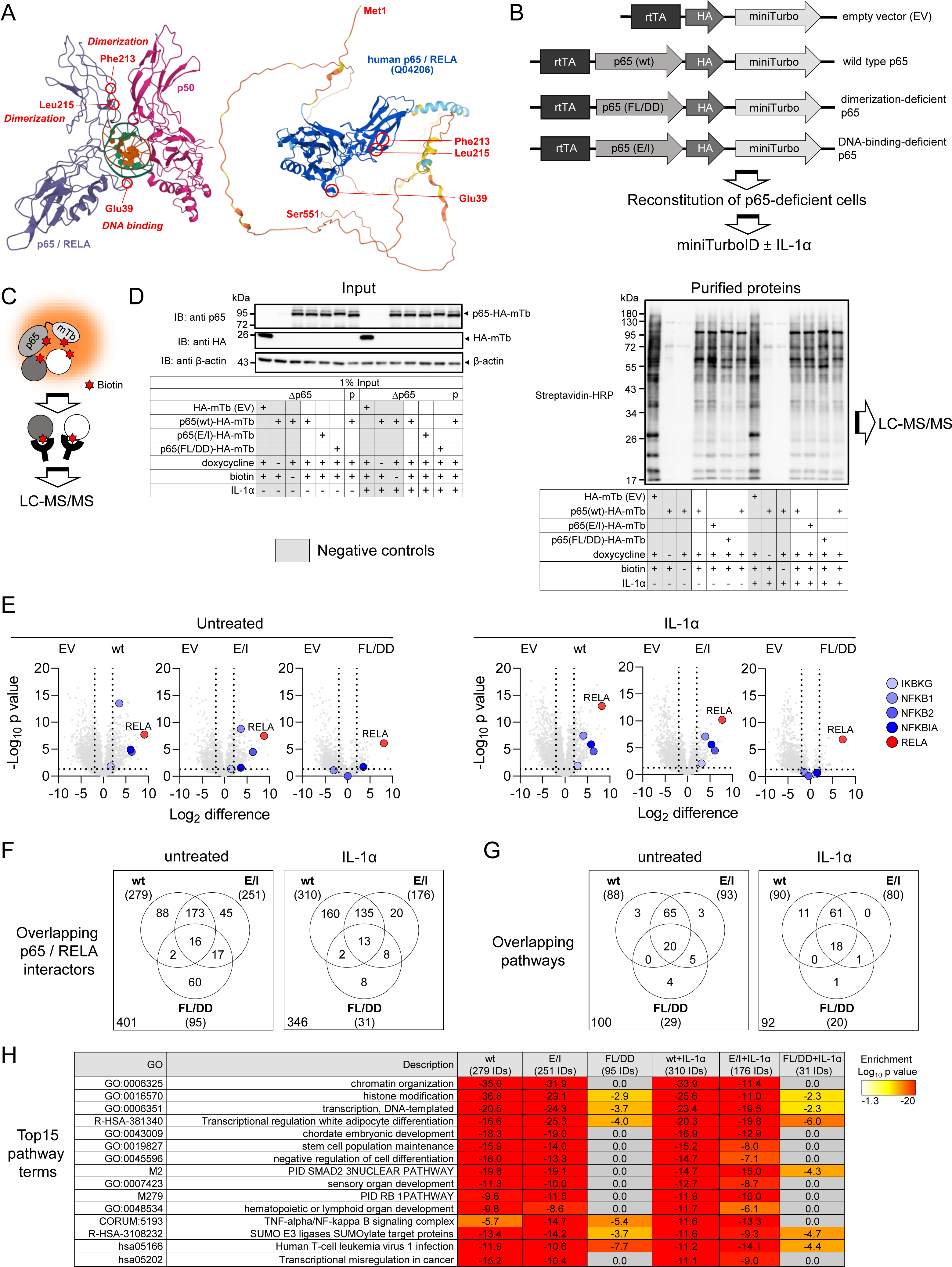
Proteome-wide identification of the dimerization- or DNA-binding dependent p65 / RELA interactomes by proximity-labeling. (A) The left graph shows the X-ray crystal structure of a p50 / p65 heterodimer bound to DNA as published in (Chen et al., 1998) (PDB 1kvx), while the right graph shows the entire p65 protein structure including the disordered C-terminal half as calculated by alphafold (https://alphafold.ebi.ac.uk/entry/Q04206). Residues required for dimerization (Phe (F) 213, Leu (L) 215) or DNA binding (Glu (E) 39) are indicated in both structures. (B) Scheme of the HA-tagged p65-miniTurbo fusion proteins that were used to reconstitute p65-deficient HeLa cells under the control of a tetracycline-sensitive promoter. F213 and L215 in p65 wildtype (wt) were mutated to Asp (FL / DD) for dimerization-deficient p65 or E39 to Ile (E / I) for DNA-binding-deficient p65. (C) Principle of proximity-based biotin tagging. (D) Pools of HeLa cells with CRISPR / Cas9-based suppression of endogenous p65 / RELA (Δp65) were transiently transfected (using branched Polyethyleneimine, PEI)) with the constructs shown in (B) and their expression was induced with doxycycline (1 µg / ml) for 17 h. At the end of this incubation, intracellular biotinylation was induced by adding 50 µM biotin for 70 minutes as indicated. Additionally, half of the samples were treated with IL-1α (10 ng / ml) for the last 60 minutes. Cell cultures expressing HA-miniTurbo only (empty vector, EV) or receiving only doxycycline or biotin served as negative controls (indicated by gray font). Parental HeLa cells (p) were included as further controls. Left panel: Cells were lysed and proteins were analyzed by Western blotting for the expression of p65-HA-miniTurbo and HA-miniTurbo using anti p65 and anti HA antibodies. Equal loading was confirmed by probing the blots with anti β-actin antibodies. Right panel: Biotinylated proteins from the same samples were purified on streptavidin agarose beads and biotinylation patterns were visualized by Western blotting using streptavidin-horseradish peroxidase (HRP) conjugates (representative images from two independent experiments). (E) Biotinylated proteins from the experiment shown in (C) and from a second biological replicate were identified by mass spectrometry. Volcano plots show the ratio distributions of Log_2_-transformed mean protein intensity values on the X-axes obtained with wild type p65 or the p65 mutants compared to the empty vector controls in the presence or absence of IL-1α treatment. Y axes show corresponding p values from t-test results. Strong enrichment of the bait p65 / RELA proteins together with the core canonical NF-kB components is shown in red and blue colors, respectively (two biologically independent experiments and three technical replicates per sample). (F) Specific proteins binding to p65 / RELA wild type were defined by significant enrichment (LFC ≥ 2, -log_10_ p ≥ 1.3) compared to HA-miniTurbo only and to cells exposed to doxycycline or biotin only (see Supplementary Fig. 2). This set of proteins was intersected with proteins enriched in cells expressing p65 mutant proteins (LFC ≥ 2, -log10 p ≥ 1.3). Venn diagrams show the numbers of p65 / RELA interactors and their overlaps before and after IL-1α-treatment, with values in the lower left corners indicating total numbers of interactors. (G) The six protein sets shown in (E) were subjected to parallel overrepresentation pathway analysis using Metascape software (Zhou et al., 2019). The Venn diagrams show the overlap of the top 100 enriched pathway terms. For IL-1α samples, only 92 terms were enriched. Values in the lower left corners indicate total numbers of unique pathways. (H) The table shows the most strongly enriched pathway categories associated with the p65 / RELA wild type or mutant interactomes. Numbers in brackets indicate the total numbers of p65 / RELA interactors per condition that were subjected to overrepresentation analysis according to (E, F). The mass spectrometry data and bioinformatics analysis results are provided in Supplementary Table 1. See also Supplementary Fig. 1 and 2. rtTA, reverse tetracycline-controlled transactivator.

To comprehensively determine the p65 / RELA interactome as it occurs in intact cells, we constructed expression vectors containing wild type p65 / RELA or the E / I and FL / DD mutants fused in frame to a C-terminal HA-tag and a modified miniTurbo biotin ligase (Branon et al., 2018) **(Fig. 1B)**. The fusion proteins were expressed under the control of a tetracycline-sensitive promoter using a single vector system **(Fig. 1B)**. Parental HeLa cells and stable HeLa cell lines genetically edited by CRISPR-Cas9 to have strongly reduced p65 / RELA levels were reconstituted with the constructs and showed doxycycline-dependent expression, nuclear translocation of NF-κB and basal as well as IL-1α-inducible transcriptional activity sensitive to the aforementioned mutations, demonstrating that the fusion proteins were functional in the NF-κB system **(Supplementary Fig. 1A-C)**.

Following intracellular expression, proteins in the vicinity of p65 / RELA were supposed to be modified with biotin in a distance-dependent manner by the miniTurbo part, as shown schematically in Fig. 1C. Wild type p65 / RELA or the two mutant versions were expressed at comparable levels **(Fig. 1D, left panel)**. Biotinylated proteins were purified from extracts of untreated cells or cells exposed to IL-1α for 1 h and visualized by streptavidin-HRP cojugates **(Fig. 1D, right panel).** Biotinylation was strictly dependent on the simultaneous addition of doxycycline and biotin to the cell cultures, as no signals were detected with doxycycline or biotin alone **(Fig. 1D, right panel)**. The latter conditions, together with samples from cells expressing HA-miniTurbo alone (EV), served as important negative controls to define specific p65 / RELA interactors in the bioinformatics analyses later on **(Fig. 1D, gray colors)**.

Across all conditions, a total of 3,928 protein IDs were identified from purified biotinylated proteins by LC-MS / MS from the two biological replicates **(Supplementary Table 1)**. Volcano plot analyses show that many of these proteins are labeled by the small and presumably more mobile HA-miniTurbo protein **(Fig. 1E, samples labeled EV)**. In contrast, p65 / RELA was highly enriched along with its canonical interaction partners p50 (NFKB1), p52 (NFKB2), IκBα (NFKBIA), and NEMO (IKBKG) in samples expressing the fusion proteins **(Fig. 1E)**. Based on a significant at least fourfold enrichment compared with all negative controls, we found 279 specific p65 / RELA interactors in untreated cells and 310 in IL-1α-treated cells **(Fig. 1E, Supplementary Fig. 2A-B)**. With the E / I mutant, 251 interactors were identified in comparison, compared with only 176 after IL-1 treatment **(Fig. 1E)**. Striking was the significantly reduced number of p65 / RELA interactors in the FL / DD mutant, which amounted to 95 in untreated cells and only 31 after IL-1 treatment **(Fig. 1F)**. Since, as shown in **Fig. 1D**, a comparable enrichment and thus (auto)biotinylation of the RELA (FL / DD) fusion protein was measured, we consider this effect to be specific. Of 401 specific interactors in untreated cells, only 16 (4%) were associated with all p65 / RELA bait proteins, and these numbers (13 or 4%) were similar after IL-1α treatment **(Fig. 1F)**.

To investigate the extent to which these differences were also reflected at the functional level, comparative overrepresentation analyses were performed for all six protein groups shown in **Fig. 1G** to identify the 100 most enriched pathways. These data show that the p65 / RELA wild type and the E / I mutant, but not the FL / DD mutant, behave largely similarly in terms of biological function of the interacting proteins **(Fig. 1G, Supplementary Table 1)**, whereas the FL / DD mutant is essentially associated with a loss of pathway terms **(Fig. 1G)**. This is particularly evident when considering the 15 most enriched pathways, which include processes such as chromatin organization, histone modifications, transcription, NF-κB signal transduction, and various developmental and differentiation steps **(Fig. 1H)**.

Next, particularly in light of the many molecular functions in gene regulation attributed to p65 / RELA (Martin et al., 2020), we focused on a detailed analysis of the composition of the p65 / RELA interactome, primarily concerning proteins with a role in chromatin-associated processes or transcription.

In total, we found 46 proteins for which an interaction with p65 / RELA had already been documented in the STRING database at different experimental levels of evidence **(Fig. 2A)**. For the most part, these factors were strongly enriched by miniTurboID and included many well-characterized transcription factors (e.g. NFATC2, IRF1, FOSL1, CEBPa/d, JUN), histone-modifying enzymes (e.g. EP300, CREBBP, KAT2A), chromatin remodelers (e.g. DPF1/2, NCOR2), nuclear cofactors (e.g. MED15, BCL6) and signaling factors (e.g. IRAK1) **(Fig. 2A)**.

**Fig. 2.**
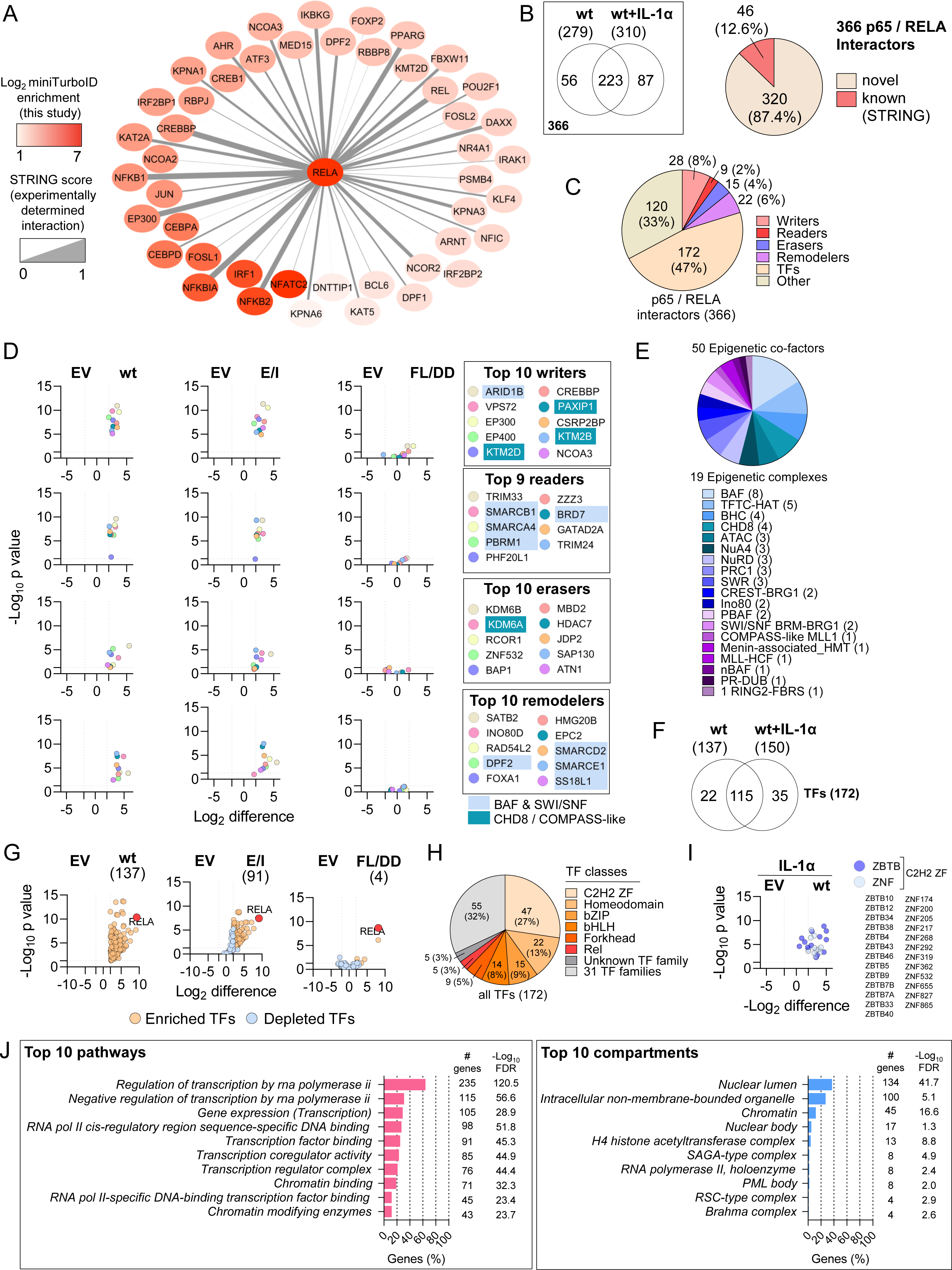
Protein composition of the p65 / RELA interactome. (A) Protein interaction network of the 46 known p65 / RELA interactors found by miniTurboID. Edge widths visualize the evidence for experimental interactions deposited in the STRING database (Szklarczyk et al., 2019). Nodes are colored in red and are arranged according to the enrichment found by proximity labeling in our study. (B) Venn diagram of p65 / RELA interactors in IL-1α or untreated cells revealing a total of 366 unique p65 / RELA interactors, of which 320 (87.4 %) have no documented protein interaction entries in STRING. (C) Overlap of the RELA interactome with 1639 human TFs (Lambert et al., 2018) and 801 epigenetic regulators (Marakulina et al., 2023). (D) Graphs visualizing the top 10 enriched epigenetic regulators. Volcano plots show the ratio distributions of Log_2_ transformed mean protein intensity values obtained with wild type p65 / RELA (wt) or with p65 / RELA mutants (FL/DD, E/I) compared to empty vector controls (EV). Only 9 reader proteins were found. (E) Association of enriched epigenetic regulators with known epigenetic complexes according to the annotation provided by (Marakulina et al., 2023). Numbers in brackets show identified components per complex. (F) Venn diagram showing the overlap of enriched TFs in basal or IL-1α-stimulated conditions. (G) Volcano plots visualizing all TFs significantly enriched with wt p65 / RELA (LFC ≥ 2, -log_10_ p ≥ 1.3) compared with empty vector control (EV) and the changes obtained with p65 mutants in basal conditions. (H) Distribution of TF families found to be associated with p65 / RELA in basal and IL-1α-stimulated conditions according to the annotation provided by (Lambert et al., 2018) (I) IL-1α-dependent enrichment of all TF belonging to ZBTB and ZNF families as identified by miniTurboID. (J) The top 10 pathway terms according to GO (BP, CC, MF), KEGG, Reactome, STRING clusters and WikiPathways data base entries and the top 10 subcellular localizations associated with the 366 p65 / RELA interactors. Annotations, number of components and false discovery rates (FDR) were retrieved using the STRING plugin of Cytoscape (Shannon et al., 2003). The mass spectrometry data sets and bioinformatics analysis results are provided in Supplementary Table 1.

Because the total number of p65 / RELA interactors in untreated and IL-1-treated cells, respectively, comprised 366 proteins, we conclude that 87% of the proximity-based p65 / RELA interactome consists of novel interactors **(Fig. 2B)**.

When compared to the list of 1639 TFs documented in the human genome by Lambert et al. (Lambert et al., 2018), 172 (47 %) of all p65 / RELA interactors are classified as DBD-containing TF proteins **(Fig. 2C)**. Based to 801 epigenetic regulators contained in the newest version of the Epifactors data base (Marakulina et al., 2023), a further 74 (20 %) of all p65 / RELA interactors are chromatin writers, readers, erasers or remodelers (**Fig. 2C**).

The interaction of p65 / RELA with these nuclear cofactors was completely abolished by the FL / DD (but not the EI) mutant in almost every case, as shown in **Fig. 2D** using the top10 strongest p65 / RELA interactors as an example. Based on annotations in the Epifactors database, 50 of the epigenetic regulators are subunits of a total of 19 established nuclear multiprotein complexes, with components of the BAF complex being most abundant in the p65 / RELA interactome (**Fig. 2E, Supplementary Table 1**). Particularly prominent were four to eight subunits each of the BAF, TFC-HAT, BHC and CHD8 complexes. A total of 9 factors were assigned to the canonical BRG / BRM-associated BAF (BAF or cBAF), polybromo-associated BAF (PBAF), non-canonical BAF (ncBAF) and mammalian SWI/SNF (short for SWItch/sucrose nonfermentable) complexes (**Fig. 2D, highlighted in light blue**). Four factors belonged to the CHD8 or COMAPSS (short for proteins associated with Set1C) complexes (**Fig.2D, highlighted in turquoise).**

Of the 172 TFs that interacted with p65 / RELA in total, 137 were already enriched under basal conditions **(Fig. 2F)**. Of this group, 91 (66 %) were still present with the E / I mutant and only 4 (3 %) with the FL / DD mutant, indicating that only about 1/3 of the TF interactions of p65 / RELA require DNA binding, whereas virtually all TF interactions require dimerization. (**Fig. 2G**). 117 of the 172 TFs found in basal or IL-1α-stimulated conditions, were distributed among 7 TF classes, with C2H2 ZF and homeodomain TFs being the most abundant and accounting for 40% of all p65 / RELA interactors (**Fig. 2H**). bZIP TFs were the third most abundant and contained a number of already known p65 / RELA interactors such as JUN, ATF2 and FOSL2, among others (**Fig. 2A and Fig. 2H**).The remaining 55 TFs represented 31 different TF classes (**Fig. 2H**). Overall, we found that in IL-1α stimulated cells, 13 ZBTB and 12 ZNF transcription factors, both from the C2H2 ZF class, were the most frequently identified p65 / RELA interactors among all enriched TF families (**Fig. 2I**).

In terms of molecular functions, the 366 p65 / RELA interactors shown in **Fig. 2B** are almost exclusively associated with RNA polymerase II-regulated transcription processes **(Fig. 2J, left panel)** and are localized largely in the nucleus, in membrane-less organelles, and in chromatin **(Fig. 2J, right panel)**.

Taken together, these data demonstrate that proximity labeling reveals a much larger p65 / RELA interactome and its dynamics than previously appreciated. Through this approach, we find a complex DNA-binding, dimerization-, and IL-1α-dependent remodeling of the p65 / RELA interactome, with mutation of only two dimerization-related amino acids in RHD exerting the strongest influence, consistent with the interpretation that most p65 / RELA interactions with other cellular proteins do not require direct or stable interactions with DNA but rely primarily on intact dimerization functions. Half of the p65 / RELA interactome is dominated by a large number of TFs distributed across many different classes, of which about one third also require an intact RELA DBD in addition to dimerization. The second largest group, besides TFs, is represented by components from protein complexes that affect chromatin modifications and remodeling and whose interaction with p65 / RELA appears to be largely independent of DNA-binding. Overall, based on bioinformatic analyses, 87% of the p65 / RELA interactors are novel, defining a previously unknown dimension of the extensive interaction of p65 / RELA with other transcriptional regulators and cofactors.

### Functional validation of 38 p65 / RELA high confidence interactors by a targeted siRNA screen

Based on greater than 8-fold enrichment in both replicates in untreated or IL-1α-stimulated cells and lack of published evidence for clearly defined functions in the NF-κB system, we finally extracted a list of 38 “high confidence” p65 / RELA interactors (HCI) **(Fig. 3A)**.

**Fig. 3.**
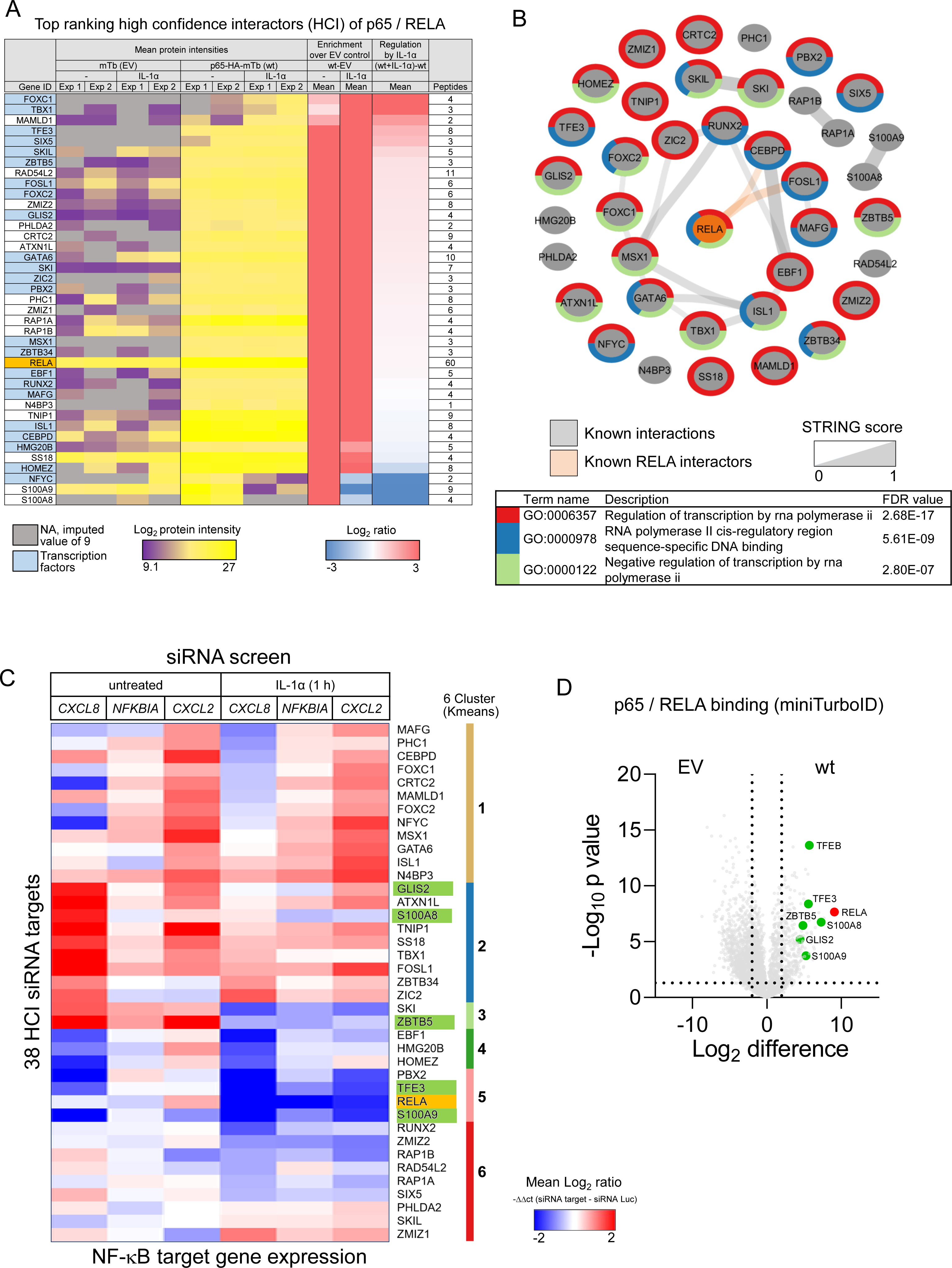
A targeted siRNA screen of 38 novel high confidence interactors (HCI) of p65 / RELA shows their function in regulation of prototypical NF-kB target genes. (A) Final list of top ranking high confidence interactors p65 / RELA selected for further studies. The heatmap shows the Log_2_ transformed mean protein intensity values from technical triplicates of the two biological independent miniTurboID experiments, the enrichment ratio values compared to the empty vector (HA-miniTurbo) control (EV) and the regulation by IL-1α. With the exception of N4BP3, all proteins were identified by at least two peptides. (B) Graph showing that the top 38 p65 / RELA interactors are largely devoid of known protein interactions based on STRING entries. According to STRING, only two factors (CEBPD and FOSL1) interact with p65 /RELA. Node borders visualize the main functional annotations. (C) HeLa cells were transiently transfected for 48 h with 20 nM of siRNAs mixtures for 38 HCI and p65 / RELA, a siRNA targeting luciferase, transfection reagent alone or were left untreated (untr.). Half of the cells per plate were treated for 1 h with IL-1α (10 ng / ml) at the end of the incubation. cDNAs were transcribed in lysates and amplicons for three NF-kB target genes, two housekeeping genes and all 38 HCI p65 / RELA interactors were pre-amplified by linear PCR and then quantified by qPCR. Based on Ct values, mRNA levels were quantified and normalized against *GUSB*. The effects of knockdowns were calculated separately for basal and IL-1α-inducible conditions against the luciferase siRNA. The heatmap shows hierarchically Kmeans clustered mean ratio values derived from three biologically independent siRNA screens. As a positive control, RELA knockdowns were performed in parallel. Green colors highlight p65 / RELA interactors selected for further analysis. (D) The miniTurboID enrichment of six p65 / RELA interactors (green colors) chosen from (C) is shown. The complete set of data of the screen is provided in Supplementary Table 2. See also Supplementary Fig. 3.

As before, these proteins were highly associated with transcriptional functions **(Fig. 3B)**. Because they had almost no known interactions with each other and only two had a documented interaction with p65 / RELA (CEBPD and FOSL1) in the STRING data base, this set defines a new, previously unknown part of the p65 / RELA interactome that we selected for follow-up validation (**Fig. 3B)**.

We then performed a targeted siRNA screen to investigate all 38 HCI concerning their relevance for NF-κB-mediated gene expression. Specifically, we used commercial siRNAs to suppress all 38 factors individually in untreated and in IL-1α-stimulated cells and then assessed the effects on the expression levels of endogenous NF-κB target genes. The screen was performed in a miniaturized format, by which RT-qPCRs were performed in total cell lysates with an intermediate linear pre-amplification PCR step, using gene-specific primers / Taqman probes for all of the targeted genes (38 plus p65 / RELA), three IL-1α-inducible genes (*CXCL8*, *NFKBIA*, *CXCL2*) and two housekeeping genes (*GUSB*, *GAPDH*) **(Supplementary Fig. 3A)**.

Results from three independent screens were normalized and expressed as mean difference between the siRNA target and a control siRNA directed against luciferase. The mRNA levels of 38 HCI were successfully suppressed by the siRNAs, while there was little effect on *GUSB* and *GAPDH* (**Supplementary Fig. 3B, Supplementary Table 2**).

As expected, the strongest suppression of IL-1α-inducible expression of *CXCL8*, *NFKBIA* and *CXCL2* was observed with p65 / RELA knockdown **(Fig. 3C)**. However, we found that for all tested genes and conditions the knockdown of a single HCI affected at least one NF-κB target gene in basal or IL-1α-stimulated conditions, or both **(Fig. 3C)**.

Hierarchical clustering of the expression patterns revealed three clusters (3-6) of HCI whose suppression resulted in relatively uniform and strong suppression of IL-1α target genes, while cluster 2 comprised a set of genes whose suppression had a strong effect on the basal expression of *CXCL8* and *CXCL2* **(Fig. 3C)**.

While p65 / RELA seemed to be essential, these data showed a functional contribution of all 38 top HCI to NF-κB-dependent gene expression and suggest that each interacting protein in a (gene-)specific manner shapes the basal and inducible state of p65 / RELA-dependent genes.

To further investigate the contribution of HCI to the regulatory functions of p65 / RELA, we selected six transcription factors for more detailed and genome-wide follow-up studies, namely zinc finger and BTB domain containing 5 (ZBTB5, encoded by KIAA0354), Zinc finger protein GLI-similar 2 (GLIS2, also called Neuronal Krueppel-like protein, NKL), S100A8 (also called CAGA, CFAG, MRP8), and S100A9 (also called CAGB, CFAG, MRP14), and transcription factors E3 (TFE3, BHLHE33) and EB (TFEB, BHLHE35) **(Fig. 3C)**. All six proteins clearly interacted with p65 / RELA in the miniTurboID screen **(Fig. 3D)**. The interactions of TFE3, TFEB, GLIS2 and ZBTB5 with p65 / RELA were also confirmed for the endogenous proteins at single cell level by proximity-ligation assays (PLA) and were significantly reduced in p65-deficient cells (**Supplementary Fig. 4).**

This selection of factors provided an opportunity to validate the p65 interactors using the example of two poorly characterized transcription factors with completely unknown relationships to p65 / RELA (ZBTB5, GLIS2) and two pairs of related factors (S100A8 / A9, TFE3 / TFEB) that play a role in inflammation but do not have a well-established mechanistic link to the NF-κB system.

### Crosstalk of lysosomal transcription factors TFE3 / TFEB and GLIS2 with the (inducible) NF-κB system

The miniTurboID data showed that 3 out of 4 MiT-TFE family members, i.e. TFE3, TFEB and microphthalmia-associated transcription factor (MITF) and all three GLIS family members (GLIS1-3) bound to p65 / RELA wt **(Fig. 4A)** This interaction was largely abolished in the FL / DD dimerization –deficient p65 / RELA mutant for all six factors and reduced in the E / I DNA-binding deficient mutant mainly after cytokine stimulation **(Fig. 4A)**.

**Fig. 4.**
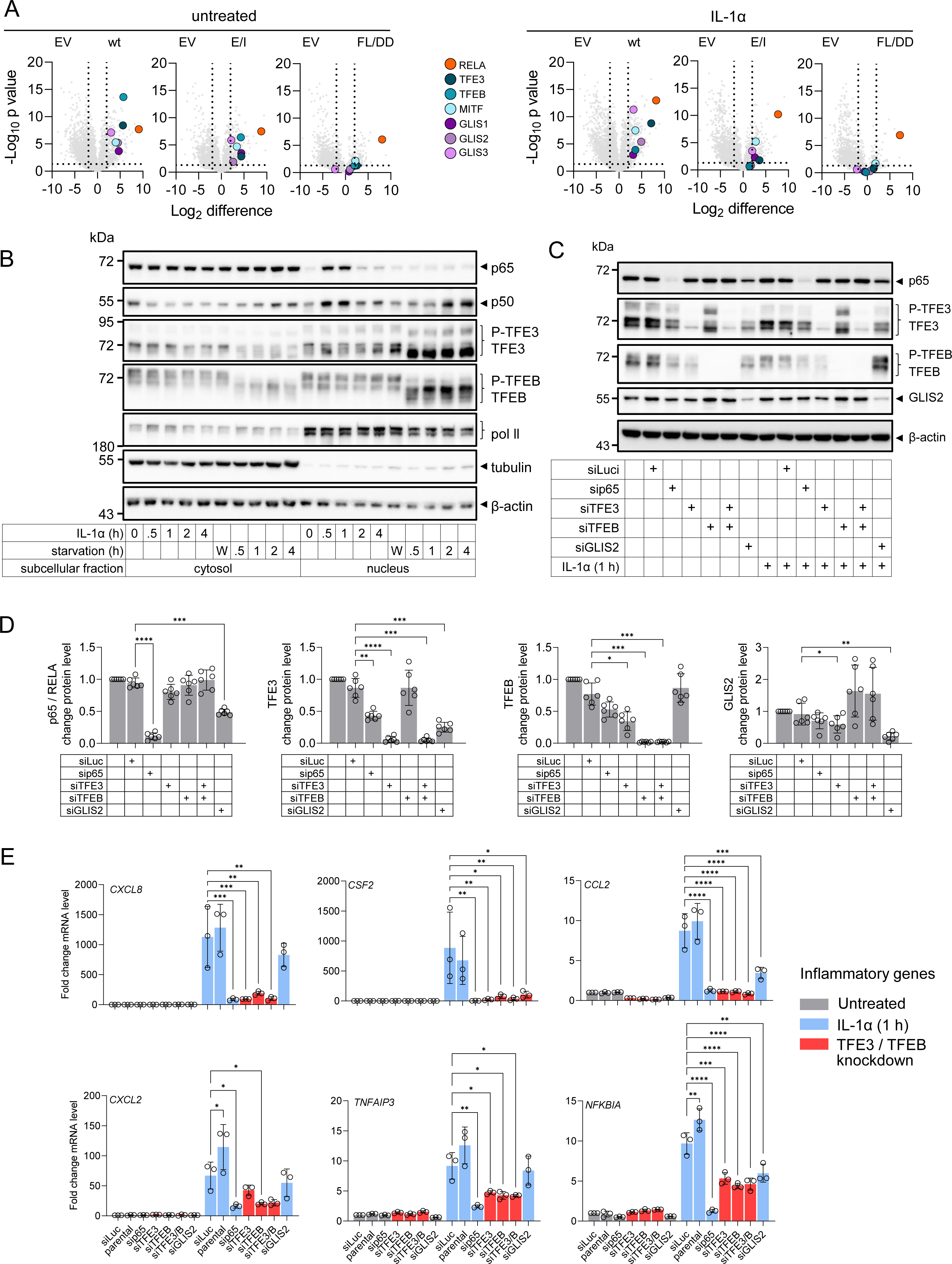
Interdependent regulation of NF-kB-target genes by TFE3, TFEB and GLIS2. (A) Volcano plots revealing the basal and IL-1α-dependent enrichment of all MiT / TFE and GLIS family members in the miniTurboID experiments. For details see Fig. 1D. (B) The subcellular distribution of phosphorylated (P) and dephosphorylated forms of TFE3 and TFEB was evaluated by Western blotting in cell extracts from HeLa cells stimulated with IL-1α or subjected to starvation (HBSS) for the indicated times. “W” indicates samples washed four times with HBSS and then supplemented with their previous cell culture medium to control for effects caused by the washing procedure prior to addition of starvation medium. Antibodies against RNA polymerase (pol II), tubulin or β-actin served as control for fractionation and equal protein loading. Shown is one out of three biologically independent experiments. See Supplementary Fig. 5 for quantification of replicates. (C) Parental HeLa cells or cells transfected with siRNAs (20 nM) against TFs or luciferase (as a negative control) were cultivated for 48 h. Then, half of the cells were stimulated for 1 h with IL-1α (10 ng / ml) or were left untreated. Total cell extracts were examined for the expression of the indicated proteins by Western blotting. Antibodies against β-actin served as loading controls. Shown is one out of three biologically independent experiments. (D) Quantification of the basal expression levels of the indicated TFs in extracts of cells transfected as in (C). Data show mean values relative to parental cells ± s.d. from six biologically independent experiments. Asterisks indicate p values (*p ≤ 0.05, **p ≤ 0.01, ***p ≤ 0.001, ****p ≤ 0.0001) obtained by two-tailed unpaired t-tests. (E) Total RNA isolated from cells treated as in (C) was analyzed for mRNA expression of the indicated NF-kB target genes by RT-qPCR. Data show mean values relative to cells transfected with luciferase siRNA ± s.d. from three biologically independent experiments. siTFE3 / B indicates double knockdown of TFE3 and TFEB. Asterisks indicate p values (*p ≤ 0.05, **p ≤ 0.01, ***p ≤ 0.001, ****p ≤ 0.0001) obtained by one-way ANOVA.

TFE3 and TFEB proteins appeared as multiple bands in cell extracts in the absence of nutritional stress and their dephosphorylated, faster-migrating forms rapidly accumulated in the nuclear fraction upon starvation as described before **(Fig. 4B and Supplementary Fig. 5)** (Martina et al., 2016; Martina et al., 2014; Settembre et al., 2011). Under non-starved conditions, considerable amounts of both phosphorylated TFE3 and TFEB were present in the nucleus **(Fig. 4B and Supplementary Fig. 5)**. IL-1α treatment did not change the phosphorylation patterns and the subcellular distributions of TFE3 / TFEB, but caused transient nuclear translocation of p65 and p50 between 0.5 and 1 h as expected **(Fig. 4B and Supplementary Fig. 5)**.

Silencing of p65 / RELA, TFE3 / TFEB and GLIS2 by siRNA revealed their profound suppression at the protein levels **(Fig. 4C)**. We also observed that p65 / RELA knockdown reduced TFE3 levels, TFE3 knockdown reduced TFEB and GLIS2 levels, and GLIS2 knockdown reduced p65 / RELA and TFE3 levels, suggesting a mutual regulation of the three factors at the protein level **(Fig. 4D)**.

RT-qPCR analysis of six prototypical TFE3 / TFEB autophagy and lysosomal target genes with a conserved cis-element in the regulatory region, the so-called Coordinated Lysosomal Expression and Regulation (CLEAR) element (Martina et al., 2016; Martina et al., 2014; Sardiello et al., 2009), revealed no effect of silencing p65 / RELA on mRNA expression of any of these genes **(Supplementary Fig. 6)**. These CLEAR genes were also not regulated by IL-1α **(Supplementary Fig. 6)**.

In contrast, knockdown of TFE3 or TFEB strongly suppressed the IL-1α-mediated expression of five prototypical inflammatory IL-1α target genes (*IL8/CXCL8*, *CSF2*, *CCL2*, *TNFAIP3*, *NFKBIA*), while GLIS2 knockdown suppressed the IL-1α-mediated expression of *CSF2*, *CCL2* and *NFKBIA* **(Fig. 4E)**.

These data provided additional functional validation of three of the p65 / RELA interactors and suggested a unidirectional crosstalk of lysosomal transcription factors TFE3 / TFEB with the IL-1α-NF-κB system.

### ZBTB5, GLIS2, S100A8 / A9 and TFE3 / TFEB are co-regulators of the p65 / RELA gene response

We then assessed genome-wide roles of the six selected factors. In a first series of transcriptome analyses using 48,000-probe microarrays, we compared silencing of p65 / RELA with silencing of ZBTB5 and S100A8 / A9, whereas in a second series we compared silencing of p65 / RELA with silencing of GLIS2, TFE3, and TFEB. Differentially expressed genes (DEGs) in the seven knockdowns were defined based on a Log_2_ fold change (LFC) ≥ 1 with a -Log_10_ p value ≥ 1.3 compared to luciferase siRNA transfections.

Each knockdown affected more than 1000 target genes in untreated cells across the entire expression range of all genes, illustrating the broad role of RELA and its HCI in homeostatic cell functions **(Supplementary Fig. 7A-B, Supplementary Table 3)**.

Next, we addressed the question of the extent to which the six factors are involved in (co)regulation of the IL-1α-regulated NF-κB response **(Fig. 5A)**.

**Fig. 5.**
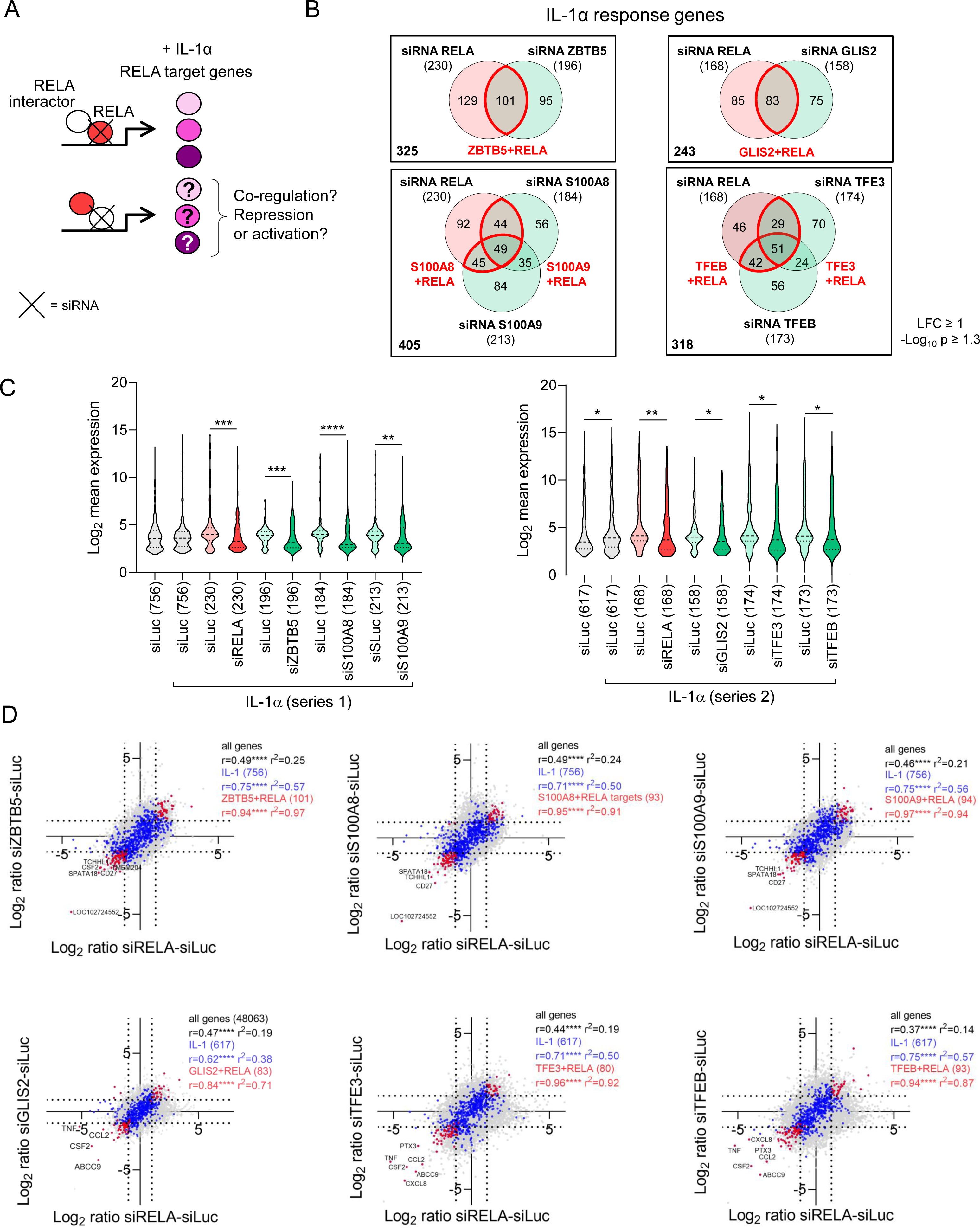
ZBTB5, GLIS2, S100A8 / S100A9 and TFE3 / TFEB co-regulate IL-1α-inducible subsets of RELA target genes. (A) Schematic illustrating the strategy to analyze the influences of novel p65 / RELA interactors on IL-1α-regulated p65 / RELA target genes by combining siRNA-mediated knockdown with transcriptome analysis. (B) HeLa cells were transiently transfected for 48 h with 20 nM siRNA mixtures against RELA, ZBTB5, S100A8, S100A9 (series 1) or RELA, GLIS2, TFE3, TFEB (series 2) and an siRNA against luciferase (siLuc) as control. Half of the cells were treated with IL-1α (10 ng/ml) for 1 hour at the end of incubation, and Agilent microarray analyses were performed from total RNA. Normalized data were used to identify DEGs based on an LFC ≥ 1 with a -log_10_ p value ≥ 1.3. Venn diagrams show the overlap of all DEGs that were affected at least twofold by siRNA knockdown in IL-1α-treated samples, with the ratio of siLuc to individual knockdown determined in each case. Red colors mark genes jointly regulated by knockdown of RELA and one of its interactors (two biologically independent experiments). (C) Violin plots show the distribution, medians, and interquartile ranges of normalized expression levels for all IL-1α-regulated genes and the corresponding changes in the gene subsets defined in Fig. 5B that were affected by siRNA knockdown. The number of these genes is indicated in parentheses. Asterisks indicate significant changes as determined by a two-tailed Mann-Whitney test (*p ≤ 0.05, **p ≤ 0.01, ***p ≤ 0.001, ****p ≤ 0.0001). (D) Superimposed pairwise correlation analyses of the mean ratio changes of all genes (gray), IL-1α-regulated genes (blue), and gene sets significantly up- or down-regulated by siRNA knockdown (red). Ratio values from RELA knockdown conditions were compared with the knockdown of a RELA interactor in each case. Genes that are jointly regulated by knockdown of RELA and one of its interactors correspond to the Venn diagrams of (B) and are marked in red. Coefficients of correlation (Pearson’s r), corresponding p values and coefficients of determination (r^2^) rare indicated for all comparisons. The complete set of data is provided in Supplementary Table 3.

Of the 756 (series 1) and 617 (series 2) IL-1α-induced genes, 230 (30%) and 168 (27%) genes, respectively, were expressed in a p65 / RELA-dependent manner **(Fig. 5B)**.

Each individual knockdown affected a comparable number of IL-1α target genes, which partially overlapped with the p65 / RELA-regulated sets of genes as indicated by red colors in the Venn diagrams shown in **Fig. 5B**.

Similar to RELA, suppression of ZBTB5, GLIS2, S100A8 / A9, and TFE3 / TFEB overall resulted in a significant reduction in expression of their respective sets of IL-1α target genes, consistent with them acting primarily as coactivators in the IL-1 system **(Fig. 5C)**.

This phenomenon was not observed for the large number of genes expressed in basal conditions, the majority of which did not overlap with RELA target genes and thus appeared to be induced or repressed by the knockdowns in comparable proportions **(Supplementary Fig. 7C, Supplementary Table 3)**.

Correlation analyses of the effects of p65 / RELA knockdowns with the respective knockdown of a p65 / RELA interactor showed a pronounced coregulation of jointly regulated IL-1α target genes, which are represented by the red sets of genes in **Fig. 5B** and **Fig. 5D**, into the same direction. This means, if a gene was suppressed or induced with the p65 / RELA siRNA relative to the luciferase siRNA, this was also the case with the knockdown of the respective interactor **(Fig. 5D)**.

This effect was also observed for the relatively small sets of genes specifically overlapping with p65 / RELA targets in basal gene expression **(Supplementary Fig. 7D, Supplementary Table 3)**.

These data emphasize the broader relevance of the new p65 / RELA interactors at the functional level, by showing that all representatively selected factors affect specific sets of RELA target genes in homeostatic conditions and profoundly participate in the regulation of inducible subsets of the IL-1α gene response.

### p65 / RELA, ZBTB5, GLIS2, S100A8 / A9, and TFE3 / TFEB engage in complex multilayer gene regulatory networks

To identify additional functional connections between all of the groups of genes deregulated by siRNA knockdowns in the IL-1α response as shown in **Fig. 5B**, we used their STRING entries of functional protein-protein interactions to construct multidimensional interaction networks.

The basic idea of this analysis is that the gene products regulated by RELA or a RELA interactor may themselves have many other direct or functional protein interactions, providing the cell with a much larger and ultimately interconnected gene regulatory network as shown schematically in **Fig. 6A**.

**Fig. 6.**
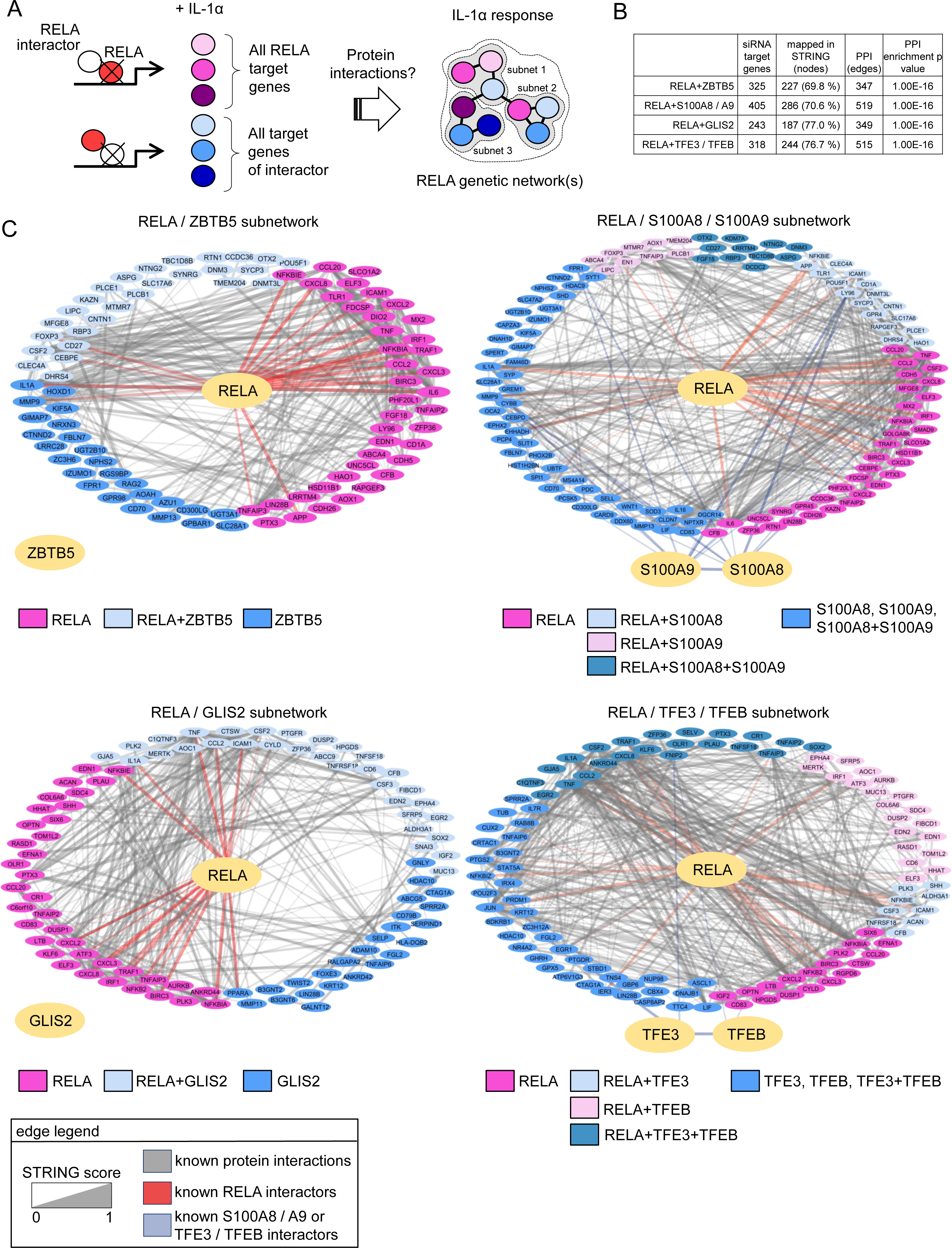
IL-1α-regulated RELA genetic networks derived from genome-wide loss of function analysis of its interaction partners. (A) Schematic illustrating the strategy to project the protein interactions of all target genes defined by knockdowns of p65 / RELA or its interactors in IL-1α-stimulated cells into combined functional networks. (B) Table summarizing the numbers of mapped IDs (= nodes) corresponding to the gene groups shown in Fig. 5B, their protein interactions (= edges) and the protein interaction network enrichment p values as derived from STRING. (C) Cytoscape-derived PPI networks. Nodes are colored and arranged according to the deregulation of the corresponding genes by knockdown of p65 / RELA or its interactors. Edges visualize known protein interactions, including the small number of interactions reported for p65 / RELA, S100A8 / 9, and TFE3 / TFEB. No interactions were found for ZBTB5 and GLIS2.

To test this hypothesis, we used the gene sets defined via Venn diagrams in **Fig. 5B** and constructed four separate PPI networks for the ZBTB5 / RELA (325 genes), S100A8 / S100A9 / RELA (405 genes), GLIS2 / RELA (243 genes), and TFE3 / TFEB / RELA (318 genes) groups. Noteworthy, only 69-77 % of the genes affected by siRNA knockdown had a PPI entry in STRING, corroborating the notion that the combined miniTurboID / transcriptome analysis effectively revealed many new components of novel genetic networks **(Fig. 6B)**.

The known interactions of factors retrieved from STRING are shown by grey lines, whereby edge width corresponds to the underlying evidence **(Fig. 6C)**. As highlighted by the light pink colors of the edges, RELA interacts with a relatively small number of proteins in all four networks. S100A8 / 9 and TFE3 / TFEB strongly interacted with each other (as expected from literature (Raben and Puertollano, 2016; Vogl et al., 2018)) and with very few other proteins, whereas ZBTB5 and GLIS2 had no known interactions at all.

The coloring of network nodes according to their dependence on RELA, ZBTB5, GLIS2, S100A8 / 9, or TFE3 / TFEB links the known levels of connectivity deposited in STRING to the novel patterns of regulation of the corresponding genes observed in our study and reveals a multitude of experimentally determined new relations between the different groups of siRNA target genes **(Fig. 6C)**.

Taken together, this refined analysis, using the IL-1 response as an example, demonstrates that RELA controls large genetic networks composed of gene regulatory subnetworks which are assembled from specific interactions of p65 / RELA with ZBTB5, GLIS2, S100A8 / 9, and TFE3 / TFEB and their target genes.

### Chromatin recruitment of RELA and its interactors

We next extended the functional analysis of selected p65 / RELA interactors to the chromatin level. For this purpose, we used our published p65 / RELA ChIPseq data from the HeLa subclone KB and screened the experimentally determined binding regions of p65 / RELA for underlying significantly enriched DNA motifs of TFE3, TFEB or GLIS2 within a range of ± 500 base pairs around the p65 / RELA peaks **(Fig. 7A, Supplementary Table 4)** (Jurida et al., 2015; Weiterer et al., 2020).

**Fig. 7.**
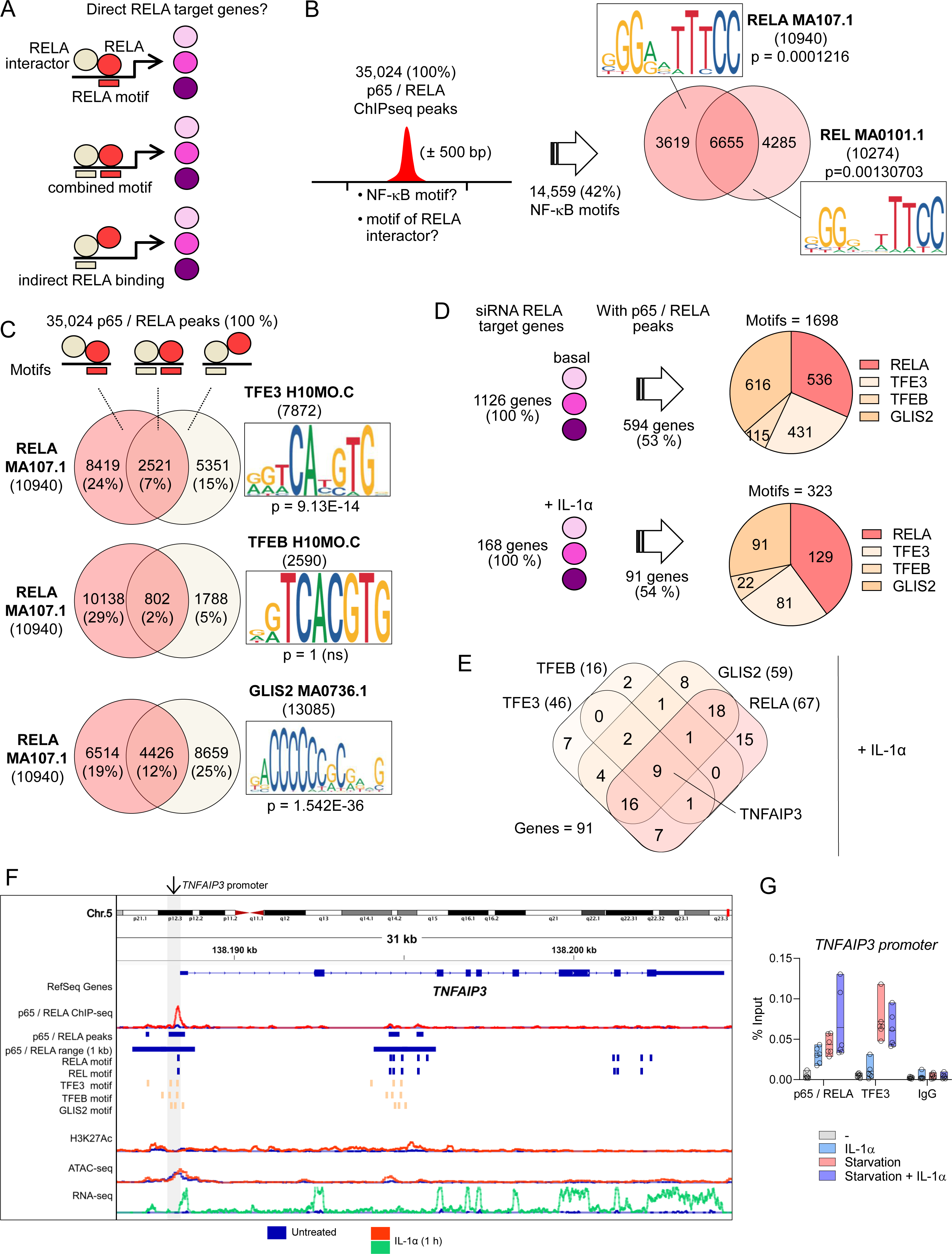
Motif analysis predicts chromatin recruitment of RELA interactors to p65 / RELA ChIPseq peaks. (A) Schematic illustrating the strategy to use p65 / RELA ChIPseq data for delineating chromatin recruitment of RELA together with its interactors on the basis of DNA motifs and three possible scenarios of interactions. (B) Windows of 1000 base pairs surrounding experimentally determined p65 / RELA ChIPseq peaks (Jurida et al., 2015) were searched for motifs of RELA and REL using matrices from the JASPAR data base. P values indicated significant enrichment compared to the whole genome. The Venn diagram shows the overlap and inserts show motif compositions. (C) Venn diagrams indicating the overlap of motifs found for RELA or the RELA interactors TFE3, TFEB or GLIS2 in chromosomal regions assigned to p65 / RELA ChIPseq peaks. P values indicated significant enrichment compared to the whole genome. Inserts show motif compositions. (D) All target genes that were significantly up- or downregulated under basal or IL-1α-stimulated conditions as shown in Fig. 5 or Supplementary Fig. 7 were collected and were examined for their association with a p65 / RELA ChIPseq peak. The pie charts show the numbers of RELA, TFE3, TFEB and GLIS2 motifs detected in siRNA RELA target genes with an annotated p65 / RELA peak in their promoters or enhancers. (E) Overlap of all genes with a p65 / RELA peak in promoters or enhancers and at least one motif for the indicated transcription factors in IL_1a-stimulated conditions. (F) Genome browser view of the *TNFAIP3* locus with p65 / RELA ChIPseq peaks, activated enhancers and promoters (H3K27ac), accessible chromatin (ATACseq) and mRNA production (RNAseq) before and after 1 h of IL-1α stimulation. Data sets were from GSE64224, GSE52470 and GSE134436 and are aligned to HG19 (Jurida et al., 2015; Weiterer et al., 2020). p65 / RELA binding regions of 1000 bp under p65 / RELA peaks and identified TF motifs are indicated by horizontal lines. (G) HeLa cells were left untreated or were starved for 24 h in HBSS. Half of the cells was treated with IL-1α (10 ng / ml) for 1 h before the end of the experiment. ChIP-qPCR was performed with the indicated antibodies or IgG controls and a primer pair covering the *TNFAIP3* promoter region (marked with an arrow in Fig. 7F). Floating bar plots show percent input plus the mean of all values from three independent biological replicates performed with two technical replicates. The complete set of data is provided in Supplementary Table 4.

First, we found that 42% of all 35,024 p65 / RELA peaks were associated with motifs specific for either p65 / RELA or for any REL (NF-κB) transcription factor (**Fig. 7B**). Second, 7 % or 12 % of all 35,024 p65 / RELA ChIPseq peaks contained a RELA motif and a TFE3 or GLIS2 motif, respectively, suggesting that RELA chromatin recruitment occurs at composite DNA binding elements which would facilitate direct binding of RELA and either TFE3 or GLIS2 to DNA (**Fig. 7C**). 24 % or 19 % of all RELA peaks contained a RELA motif, but no motif for TFE3 or GLIS2 (**Fig. 7C**). Vice versa, 15 % or 25 % of all RELA ChIPseq peaks contained a motif for TFE3 or GLIS2, respectively, but no RELA motif, suggesting indirect recruitment to DNA by PPI (**Fig. 7C**). As indicated by p values, TFE3 and GLIS2 motifs under p65 / RELA peaks were highly significantly overrepresented, compared to their distribution across the whole genome sequence **(Fig. 7C)**. This was not the case for TFEB motifs, of which 2% were found together with RELA motifs and 5% without RELA motifs at p65 / RELA peaks (**Fig. 7C**).

Around 50% of all genes which were deregulated by RELA siRNAs in basal or IL-1α-stimulated conditions (**as shown in Fig. 5 and Supplementary Fig. 7**) were associated with at least one motif for either RELA, TFE3 / TFEB, GLIS2 (**Fig. 7D**). In most instances, gene sets contained 1-3 motifs alone or in combination, in line with the notion of RELA genetic subnetworks as described before **(Fig. 6C)**. Only 9 genes were annotated with all four motifs, such as TNFAIP3 (**Fig. 7E**). The highly IL-1α-inducible TNFAIP3 gene locus contains two major p65 / RELA peaks within the promoter region that are associated with multiple TFE3, TFEB and GLIS bindings sites **(Fig. 7F)**. As a proof of concept, we chose this gene to demonstrate the IL-1α-inducible recruitment of p65 and TFE3 to the promoter of TNFAIP3 (**Fig. 7G**). Notably, this recruitment was increased by long-term starvation for 24 h, the condition known to increase translocation of TFE3 to the nucleus (**Fig. 7G**).

Similar results were obtained from the analyses of available motifs for various ZBTB family members that, in addition to ZBTB5, were identified in our p65 / RELA interactomes as shown in **Fig. 2H**. While the DNA-binding motif for ZBTB5 is unknown, ZBTB33, ZBTBT7A and ZBTB7B motifs were clearly enriched under p65 / RELA ChIPseq peaks (**Supplementary Fig. 8**). Interestingly, ZBTB7A is the only ZBTB factor, for which a role in the NF-κB system has been described. It was found to bind to p65 / RELA and control the accessibility of promoters for p65 / RELA (Ramos Pittol et al., 2018).

These combined experimental and bioinformatics analyses suggest that a considerable part of p65 / RELA chromatin recruitment could occur indirectly, through interactions with one of its many TF binding partners as defined in this study by proximity labeling.

## Discussion

Transcription factors, which account for approximately 8% of all human genes, are defined by their ability to interact with DNA and stimulate or repress gene transcription, but their experimentally determined binding sites are not necessarily predictors of the genes that they actively regulate (Cusanovich et al., 2014; Lambert et al., 2018). It has been suggested that this relative lack of specificity is compensated for by cooperativity and synergy between TFs and by their interactions with other nuclear proteins. However, understanding of these interactions and relationships is still very limited (Lambert et al., 2018). Here, we used proximity labeling to investigate the interactome of the REL family member p65 / RELA at high resolution. The data reveal hundreds of p65 / RELA interactors in a single cell type, demonstrating the enormous extent of intermolecular connectivity of a single mammalian TF. Taken together with the exemplary functional study of selected TF partners, the results have far-reaching implications for interpreting p65 / RELA-driven processes in homeostatic and diseased states.

Limited evidence exists for p65 / RELA interactors (or any other NF-κB protein) from large-scale studies. Bouwmeester et al. reported 92 p65 / RELA interactors in HEK293 cells using tandem affinity purification / mass spectrometry (TAP-MS) (Bouwmeester et al., 2004) (**Supplementary Table 5)**. By TAP-MS 2,156 high-confidence protein-protein interactions were identified from soluble and chromatin-associated complexes of 56 TFs, including 71 unique interactions for NFKB1 in HEK293 cells (Li et al., 2015) (**Supplementary Table 5)**. Goos et al. recently surveyed the PPIs of 109 human TFs using BirA fusions in HEK293 cells, finding an average of 61.5 PPIs per TF. Notably, they identified only 16 interactors for NFKB1 (p50) (Goos et al., 2022) (**Supplementary Table 5)**. The Gilmore group lists 115 RELA-interacting proteins derived from various models and systems (https://www.bu.edu/nf-kb/physiological-mediators/interacting-proteins/) (**Supplementary Table 5**), while, as shown in Fig. 2, the STRING data base currently documents less than 50 p65 / RELA interactors (Szklarczyk et al., 2019). Thus, our results significantly exceed the number of reported p65 / RELA interactors and provide in depth functional validation, which is lacking in the large scale screens cited above.

About one fifth of the p65 / RELA interactome consisted of chromatin regulators. Our data reproduced interactions of p65 / RELA with the histone acetyltransferases (HATs) p300 / CBP (also called EP or CREBBP), TIP60 (KAT5) and the histone deacetylases HDAC1 / 2 (Ashburner et al., 2001; Brockmann et al., 1999; Merika et al., 1998; Perkins et al., 1997), which were later also confirmed in the chromatin text, e.g. at H3K27-marked enhancers and promoters (Garber et al., 2012; Kim et al., 2012; Mukherjee et al., 2013; Raisner et al., 2018).

Beyond this, miniTurboID greatly advances our understanding of the complexity of the p65 / RELA nuclear cofactor interactome as with this method we detected more than 50 subunits of chromatin complexes associated with p65 / RELA. The data suggest that p65 / RELA preferentially interacts with complexes that promote active gene expression and counteract repressive programs mediated by Polycomb proteins, such as COMPASS, SWI/SNF or BAF (Cenik and Shilatifard, 2021) (Kadoch and Crabtree, 2015; Mashtalir et al., 2020) (Hodges et al., 2016; Varga et al., 2021) (Schick et al., 2021). Further p65 / RELA interactors (KDM6A, KMT2D, NCOA6, PAGR1, PAXIP1) are subunits of COMPASS-like or CHD8 complexes that regulate H3K4 methylation or chromatin remodeling at promoters and enhancers during transcriptional activation (Manning and Yusufzai, 2017; Schuettengruber et al., 2017). However, how different subunits of these complexes are assembled and recruited to chromatin for stimulus-specific functions remains an open question. Our data shed light on a possible role of the p65 / RELA interactome to coordinate these events.

In this context, it was remarkable that half of the p65 / RELA interactors represented other TFs. This result supported earlier studies showing p65 / RELA’s interaction with basic leucine zipper domain (bZIP) TFs (e.g., JUN, ATF1/2/3/7, FOSL1/2, CEBPA/D) or IRF1 (Jurida et al., 2015; Merika et al., 1998; Stein et al., 1993; Wolter et al., 2008). The sheer number of possible TF interactions suggested an extraordinary level of p65 / RELA cooperativity with this class of proteins.

NF-κB subunits are known to dimerize, potentially contributing to transcriptional selectivity (Saccani et al., 2003; Siggers et al., 2012; Smale, 2012). Testing two p65 / RELA mutants that disrupt chromatin recruitment and activation of TNFα response genes to a similar extent (Riedlinger et al., 2019), we found that dimerization played a larger role in interactions with epigenetic regulators and TFs than DNA binding. This behavior can be reconciled with the observation that, in living cells, promoter-bound NF-

κB exists in dynamic, oscillating equilibrium with nucleoplasmic dimers, with short residence times at high-affinity DNA binding sites (Bosisio et al., 2006). We suggest that miniTurboID, being crosslink-free and rapid, appears to provide a snapshot of the consequences of this dynamic equilibrium. The distinct interactomes of the E / I or the FL / DD mutants imply that, on average in the cell population studied, the majority of the p65 / RELA interactome does not require stable interactions with DNA and the complexes are (pre)assembled outside the chromatin, presumably in the nucleoplasm. These interactions are determined by dimerization properties of p65 / RELA which were diminished by FL / DD mutations. Proximity-based labeling thus informs on additional layers of TF cooperativity beyond the coordinated formation of p65 / RELA complexes on accessible chromatin templates. Such a behavior is unlikely to be captured by crosslinking-dependent techniques such as ChIPseq.

From a genome-wide perspective, the miniTurboID-based interactome suggested a model where p65 / RELA, in cooperation with its TF partners and associated epigenetic regulator complexes, instructs the cell to execute specific transcriptional programs. To test this hypothesis, we extended our analysis to three functional levels: (i) a targeted siRNA screen of a panel of high confidence interactors, (ii) a detailed identification of overlapping sets of target genes and (iii) the analysis of TF motifs under p65 / RELA peaks.

RNAi-mediated suppression of the 38 most enriched novel p65 / RELA interactors, demonstrated the gene-specific, functional contributions of 24 TFs, spanning multiple families, in regulating three canonical NF-κB target genes. These TFs exhibit varying quantitative contributions to basal and IL-1α-inducible gene expression, highlighting TF cooperativity in fine-tuning NF-κB responses.

For genome-wide loss-of-function analyses, we focused on candidates from the C2H2 (GLIS2, ZBTB5) and bHLH (TFE3, TFEB) TF families, and S100A8 / A9 as non-TF interactors with p65/RELA. GLIS2 and ZBTB5, poorly characterized TFs, are implicated in processes like epithelial-mesenchymal transition and nephronophthisis but not in the NF-κB system (Attanasio et al., 2007; Cheng et al., 2021b; de Dieuleveult and Miotto, 2018; Wilson et al., 2021). S100A8 / A9 are typical, secreted drivers of the innate immune responses, but with no role in p65 / RELA-mediated transcription (La Spina et al., 2020; Pruenster et al., 2016; Wang et al., 2018). TFEB and TFE3 are known for lysosomal gene regulation under conditions of starvation (Tan et al., 2022; Yang and Wang, 2021), but have also been suggested to contribute to LPS-mediated inflammatory gene secretion in macrophages by unknown mechanisms (La Spina et al., 2020; Pastore et al., 2016). Here, we found that the constitutively phosphorylated forms of TFE3 and TFEB contributed to IL-1α-NF-κB-regulated inflammatory gene expression in non-starved conditions. Additional data at the protein level suggest mutual regulation between p65 / RELA, TFE3 / TFEB, and GLIS2.

By intersecting transcriptome-wide analyses of cells with reduced TF levels, we identified gene sets co-regulated by p65 / RELA and six of its interactors, differing in basal conditions compared to IL-1α-activated cells. The combinatorial actions of multiple TFs and associated epigenetic regulators, as indicated by the p65 / RELA interactome, are reminiscent of gene-regulatory networks (GRN) (Spitz and Furlong, 2012). GRNs can consist of several subnetworks, each of which executes individual segments of a complex biological process (Davidson, 2010; Peter and Davidson, 2011). We tested this concept for the p65 / RELA-driven IL-1α-response by bioinformatics analyses to find that, at the systems level, a large part of the genes affected by RNAi form subnetworks with dense functional interactions. Genome-wide ChIPseq data revealed multiple motifs for TFE3 / TFEB and GLIS2 coinciding with 35,000 p65 / RELA peaks. A recent study reported a similar scale of p65 binding events across different mammalian species (31,602 to 90,570 p65/RELA peaks) (Alizada et al., 2021). NF-κB binds to both accessible and nucleosome-occluded chromatin in a TNFa-dependent manner, with defined chromatin states regulated by p65 / RELA that distinguish conserved and constitutive functions from specific pro-inflammatory functions of NF-κB (Alizada et al., 2021). Our data suggest that these chromatin states may, at least in part, be established by basal or IL-1α-regulated multi protein complexes associated with p65 / RELA.

Identifying p65 / RELA genetic subnetworks driven by p65 / RELA and its TF interactors provides a novel explanation for the NF-κB response’s stimulus-specificity and its impact on the epigenome which, in macrophages, was recently attributed to fluctuations in cellular NF-κB activity (Adelaja et al., 2021; Cheng et al., 2021a). Combining proximity-based p65 / RELA interactomics with RNAi experiments, as demonstrated in our study, thus complements and extends approaches that sought to combine ChIPseq with RNAseq to unveil the regulation of context-dependent NF-κB target genes by the two p65 / RELA TAD domains (Ngo et al., 2020).

The p65 / RELA subunit is the only NF-κB subunit whose deletion leads to embryonic lethality in mice (Beg and Baltimore, 1996). In most cells, it comprises the predominant NF-κB transcriptional activity and regulates a plethora of processes during development, the immune response and cancer (Mitchell and Carmody, 2018; Zhang et al., 2017). Consequently, the p65 / RELA pathway is tightly regulated to adjust nuclear p65 / RELA concentration and kinetics (Meier-Soelch et al., 2021). Despite significant mechanistic insights from genetically altered cells or organisms with altered p65 / RELA expression and improved knowledge of p65 / RELA-driven genetic changes, our understanding of this pathway has not yet advanced enough to yield specific and effective anti-p65 / RELA drugs. In line with a recent evaluation of NF-κB component copy numbers in various immune cells (Kok et al., 2021), our study demonstrates that quantitative proteomics can significantly contribute to understanding the NF-κB pathway’s connectivity. Assuming that stable, physically connected protein complexes are more often labeled with biotin compared to transient or indirect interactions, proximity-labeling also informs on interaction strength and frequency, which will allow further classification of p65 / RELA networks.

In summary, the high resolution novel p65 / RELA interactome and its gene-regulatory logics reported in this study provide a rich resource and a new framework for explaining p65 / RELA function in living cells.

### Limitations

Our study was restricted to a single cell type in order to standardize and integrate the different levels of molecular analyses. HeLa cells express all core components of the NF-κB pathway and have a fully functional IL-1 system (Weiterer et al., 2020). They represent one of the most widely used cellular models to date (Adey et al., 2013). Thus, we expect that p65 / RELA interactomes of other cell types will show some overlap but will also differ. A further limitation is the necessity to ectopically express a p65 / RELA fusion protein that, although performed in a p65 / RELA-deficient background using a conditional system, could influence the stoichiometry of some of the interactions we discovered. Proximity labeling generates a cumulative snapshot of protein-protein interactions, but does not allow inference of physical interactions. Because the analyses were performed on whole cell extracts, no information is available on the subcellular regulation of the interactions, although bioinformatics analyses revealed a clear predominance of nuclear p65 / RELA interactors. Despite the short labeling times (70 minutes of biotinylation), miniTurboID does not resolve the time wise sequence of interactions and rather reports cumulatively on all interactions that happened within this period of time. The generation of a specified list of 366 RELA / p65 interactors resulted from the adoption of relatively rigorously defined filtering criteria based on a combination of negative controls, levels of enrichment and T-test criteria. The same applied to the definition of p65 / RELA target genes. As with any bioinformatics analysis, it is clear that weakening or tightening these filtering criteria would increase or decrease the number of p65 / RELA interactors or genes affected by siRNA knockdown. Similarly, the annotation of factors to gene ontologies or protein networks depends on the underlying databases and the parameters set. Here, we chose to use the default settings of current versions of Metascape and STRING, respectively. However, publication of the entire raw dataset and all source data will allow colleagues in the field to (i) track all analysis results and (ii) generate alternative p65 / RELA interactor lists and GRN under their own chosen criteria.

## Methods

### Cell lines, cytokine treatment, and starvation

HeLa and KB cells were maintained in Dulbecco’s modified Eagle’s medium (DMEM; PAN Biotech; #P04-03550), complemented with 10% filtrated bovine serum (FBS Good Forte; PAN Biotech; #P40-47500) or tetracycline-free FBS (PAN Biotech; #P30-3602), 2 mM L-glutamine, 100 U/ml penicillin and 100 µg/ml streptomycin. Cells were tested for mycoplasma with PCR Mycoplasma Test Kit (Applichem; #A3744) and their identity was confirmed by commercial STR testing at the DSMZ-German Collection of Microorganisms and Cell Cultures; https://www.dsmz.de/dsmz). Stable pools of p65-depleted cells (HeLa Δp65), generated by transfections of pX459-based CRISPR/Cas9 constructs (Weiterer et al., 2020), were selected and maintained in puromycin (1 µg/ml). Prior to all experiments, puromycin was omitted for 24 h. Human recombinant IL-1α was prepared in our laboratory as described (Rzeczkowski et al., 2011) and used at 10 ng/ml final concentration in all experiments by adding to the cell culture medium for the indicated time points. Starvation of cells was induced by washing the cells four times with Hanks’ Balanced Salt solution (HBSS; PAN Biotech; #P04-32505) for the indicated time points. Starved cells were compared to non-treated control cells or cells washed four times with HBSS and then supplemented with their own culture medium to exclude effects caused by the washing procedure.

### Reagents and antibodies

Leupeptin hemisulfate (Carl Roth, #CN33.2; solved in ddH_2_O), microcystin (Enzo Life Sciences, #ALX-350-012-M001; solved in EtOH), pepstatin A (Applichem, #A2205; solved in EtOH), PMSF (Sigma-Aldrich, #P-7626; solved in EtOH), protease inhibitor cocktail tablet (Roche; #11873580001; solved in ddH_2_O), DTT (Serva; #20710.04; solved in ddH_2_O), E-64 (Sigma-Aldrich; #324890), doxycycline (Sigma-Aldrich; # D9891; solved in ddH_2_O), puromycin (Merck Millipore; #540411; solved in ddH_2_O), Biotin (Sigma-Aldrich; #B4501; solved in DMEM as 20x stock solution and sterile filtrated). Primary antibodies against the following proteins or peptides were used: Anti-β-actin (Santa Cruz; #sc-4778), anti-HA (Roche; #11583816001), anti-Phospho-IκBα Ser32 (Cell Signaling; #2859), anti-IκBα (Cell Signaling; #9242), anti-NF-κB p65 (Santa Cruz; #sc-372; #sc-8008; Bethyl Lab.; #A303-945A), anti-NF-κB p50 (Santa Cruz; #sc-8414), anti-TFE3 (Sigma-Aldrich; #HPA023881), anti-TFEB (Cell Signaling; #4240), anti-GLIS2 (Invitrogen; #PA5-40314), anti-RNA-Pol II (Millipore #17-620), anti-tubulin (Santa Cruz #sc-8035), anti-ZBTB5 (Sigma; #HPA021521), normal rabbit IgG (Cell Signaling #2729). Secondary antibodies: Dylight 488-coupled anti-mouse IgG (ImmunoReagent, #DkxMu-003D488NHSX), HRP-coupled anti-mouse IgG (DakoCytomation, #P0447), HRP-coupled anti-rabbit IgG (DakoCytomation, #P0448), HRP-Streptavidin (PerkinElmer; #NEL750001EA).

### Cloning of pTet-on-Puro-HA-miniTurbo plasmids

For generating pTet-on-Puro-HA-miniTurbo (EV, empty vector), the linker-<u>HA-miniTurbo</u> sequence was synthesized by General Biosystems and provided in donor vector pUC57-Bsal-Free. To obtain the linker-HA-miniTurbo insert, the donor vector and the target plasmid <u>pTet-on-Puro</u>-Myc-BirA were subjected to a restriction digestion reaction with FastDigest Mlul and BshTI (Agel) and subsequently ligated by T4 DNA ligase. The PCR for cloning p65 gene variants into the pTet-on-Puro-HA-miniTurbo (EV) vector was based on the donor vectors pEF-Puro-hu <u>p65 WT</u>-HA, <u>p65 E/I</u>-HA, and <u>p65 FL/DD</u>-HA (Riedlinger et al., 2019). Phusion™ high-fidelity DNA polymerase (Thermo Fisher Scientific; #F-530XL) generated the amplicons with restriction site overhangs (BshTI and Bsp1407I) and the 3-step PCR program (98°C for 30 sec, 35 x (98°C for 10 sec, 69°C for 30 sec, 72°C for 25 sec) followed by 72°C for 10 min, 4°C hold). The resulting PCR amplicons as well as the target vector pTet-on-Puro-HA-miniTurbo (EV) were digested by FastDigest BshTI (Agel) and Bsp1407I (BsrGI) and ligated. All PCR or vector digestion products were purified using the NucleoSpin® Gel and PCR Clean-Up Kit (Macherey-Nagel; #740609.50). The final plasmids were transformed into competent *E.coli XL1-Blue*, extracted by using the NucleoSpin® Plasmid or NucleoBond® Xtra Midi Kit (Macherey-Nagel; #740588.250 and #740410.50) and controlled by Sanger Sequencing (Eurofins Genomics or Seqlab Microsynth).

### Transfection of cells with branched Polyethyleneimine (PEI)

We optimized transfection conditions of the miniTurbo constructs as follows. For transient transfection of cells with expression vectors, branched polyethyleneimine (PEI; Sigma-Aldrich; #408727) was used. For T145 cell culture dishes 1 ml of pre-warmed Opti-MEM™ (serum-reduced medium; Gibco; #31985070) was mixed with 50 µg Plasmid-DNA and 120 µl ice-cold branched PEI (1 mg/ml, pH 7.0), vortexed, and incubated for 10 min at room temperature. DMEM supplemented with 10% FBS or Tet-free FBS (w/o Pen./Strep.) was added to the mixture (filled-up to 20 ml), vortexed and carefully spread over the cell layer (∼70% confluency) after the culture medium was aspirated. Cells were further incubated overnight (24 h). For other cell culture dish sizes, the volumes were adjusted accordingly.

### miniTurboID proximity labeling and purification

For each experimental condition, 5×10^5^ parental HeLa and HeLa Δp65 cells were seeded in a T145 cell culture dish and grown for 4 days. On day three, cells were transfected with pTet-On-Puro expression vectors encoding HA-miniTurbo (EV), p65(wt)-HA-miniTurbo, p65(E/I)-HA-miniTurbo or p65(FL/DD)-HA-miniTurbo using branched PEI. Following transfection, cDNA expression was induced with 1 µg/ml doxycycline for 17 h. On the next day (24 h post transfection), medium was supplemented with 50 µM exogenous biotin (Sigma-Aldrich; #B4501) 10 min prior to IL-1α treatment (10 ng/ml) for 1 h enabling biotinylation of p65 interacting proteins during inflammatory cytokine treatment. After a total of 17 h of treatment with doxycycline, cells were washed with PBS and harvested on ice by scraping and centrifugation (900 x g/4°C/5 min). Cell pellets were resuspended in 475 μl Tris/HCl (50 mM; pH 7.5) and 50 μl Triton X-100 (10% w/v). Cells were lysed by addition of 250 μl lysis buffer (50 mM Tris/HCl, 500 mM NaCl, 2% w/v SDS, add freshly 1 mM DTT, 1x Roche inhibitor cocktail) followed by incubation on ice for 10 min. Cells were sonicated (settings: 3 – 4 x 30 sec on /30 sec off, 4°C, power high; Bioruptor, Diagenode) and lysates were cleared by centrifugation at 16,000 x g at 4°C for 15 min. For validation of cDNA expression and induced biotinylation 1% of the pulldown input of the lysates (7 µl) were analyzed by SDS-Page and immunoblotting. For the pulldown of biotinylated proteins 700 μl of the lysate was added to 60 μl of streptavidin-agarose beads (Thermo Scientific; #20349) equlibrated in lysis buffer and rotated end over end overnight (16 – 18 h) at 4°C. Beads were collected by centrifugation at 1,000 x g for 2 min and were washed once with 0.5 ml wash buffer I (2% w/v SDS), twice with 0.5 ml wash buffer II (50 mM HEPES, 0.5 M NaCl, 1 mM EDTA, 0.1% w/v sodium deoxycholate, 1% v/v Triton X-100; pH 7.5), twice with 0.5 ml wash buffer III (10 mM Tris/HCl, 1 mM EDTA, 250 mM LiCl, 0.5% w/v sodium deoxycholate, 0.5% w/v NP-40; pH 7.4), twice with 0.5 ml wash buffer IV (50 mM Tris/HCl, 50 mM NaCl, 0.1% v/v NP-40; pH 7.4) and once with 0.5 ml wash buffer V (50 mM Tris/HCl; pH 7.4). The beads were resuspended in 1 ml buffer V, of which 80% were used for mass spectrometry analysis. For the validation of affinity purifications, the remaining 20% of the beads (supplemented with 40 µl of 2X ROTI®Load and boiled at 95°C for 10 min) were subjected to SDS-PAGE for immunoblotting together with 1% input samples using HRP-streptavidin conjugate (PerkinElmer; #NEL750001EA) or anti-p65 (Santa Cruz Biotechnology; #sc-8008), anti-HA (Roche; #11583816001), anti-β-actin (Santa Cruz Biotechnology; #sc-4778).

### Mass spectrometry analysis of miniTurboID

Samples bound to streptavidin-agarose beads were washed three times with 100 μl 0.1 M ammonium bicarbonate solution. Proteins were digested “on-bead” by the addition of sequencing grade modified trypsin (Serva) and incubated at 37 °C for 45 min. Subsequently, the supernatant was transferred to fresh tubes and incubated at 37°C overnight. Peptides were desalted and concentrated using Chromabond C18WP spin columns (Macherey-Nagel; #730522). Finally, Peptides were dissolved in 25 μl of water with 5% acetonitrile and 0.1% formic acid. The mass spectrometric analysis of the samples was performed using a timsTOF Pro mass spectrometer (Bruker Daltonics). A nanoElute HPLC system (Bruker Daltonics), equipped with an Aurora C18 RP column (25 cm x 75 μm) filled with 1.7 μm beads (IonOpticks) was connected online to the mass spectrometer. A portion of approximately 200 ng of peptides corresponding to 2 μl was injected directly on the separation column. Sample Loading was performed at a constant pressure of 800 bar. Separation of the tryptic peptides was achieved at 50°C column temperature with the following gradient of water/0.1% formic acid (solvent A) and acetonitrile/0.1% formic acid (solvent B) at a flow rate of 400 nl/min: Linear increase from 2% B to 17% B within 60 minutes, followed by a linear gradient to 25% B within 30 min and linear increase to 37% B in additional 10 min. Finally B was increased to 95% within 10 min and held for additional 10 min. The built-in “DDA PASEF-standard_1.1sec_cycletime” method developed by Bruker Daltonics was used for mass spectrometric measurement. Data analysis was performed using MaxQuant (version 1.6.17.0) with the Andromeda search engine and and all amino acid sequences of the Uniprot database (Uniprot Human reviewed proteins, database version 2021_03) were used for annotating and assigning protein identifiers (Tyanova et al., 2016a). Perseus software (version 1.6.14) was used for further analyses of protein intensity values (Tyanova et al., 2016b). For calculation of ratio values between conditions, biological and technical replicates from each condition were assigned to one analysis group using tools for categorical annotation rows. All values were log_2_-transformed and missing values were imputed using a log_2_ intensity value of 9, which was below the lowest intensity value measured across all samples. No further normalization of the pulldown experiments was performed to preserve the anticipated differences between samples. Enriched proteins between pair-wise comparisons were identified by Student’s T-tests using Perseus functions and were visualized by Volcano plots. Interesting groups of enriched p65 / RELA interactors were defined by log_2_ fold change (LFC) and statistical significance of changes based on -Log_10_ p values ≥ 1.3 as indicated in the legends. Subsequent filtering steps and heatmap visualizations were performed in Excel 2016 according to the criteria described in the figure legends. Venn diagrams were created with tools provided at http://bioinformatics.psb.ugent.be/webtools/Venn/. Overrepresentation analyses of gene sets were done using the majority protein IDs or gene IDs of differentially enriched proteins or mRNAs uploaded to Metascape software and processed with the predefined express settings (Zhou et al., 2019). Protein networks were inferred from filtered gene ID lists using information of the most current version of the STRING data base (https://string-db.org/) and networks were visualized and annotated with enriched pathway terms using Cytoscape, version 3.8.0 or higher and the STRING plugin (Shannon et al., 2003; Szklarczyk et al., 2019).

### Transfection of cells with siRNAs

Cells were seeded in 60 mm cell culture dishes and grown overnight. The medium was reduced to 3 ml and a transfection mixture was prepared as follows: 187.5 µl Opti-MEM™, 15 µl Hi-PerFect Transfection Reagent (Qiagen; #301705) and 60 µl of siRNA mixture (1 µM, finally 20 nM). The reaction mixture was vortexed and incubated for 10 min at room temperature, subsequently dripped on the culture dish, and gently mixed. After 6 h of incubation, the transfection mixture was aspirated and replaced by 4 ml fresh complete DMEM medium. Cells were then incubated for 48 h until they were further processed. The following FlexiTube GeneSolution siRNAs (Qiagen) were used: RELA/p65 (#GS5970), TFE3 (#GS7030), TFEB (#GS7942), GLIS2 (#GS84662), ZBTB5 (#GS9925), S100A8 (#GS6279), S100A9 (#GS6280). As a non-targeted control siRNA against Firefly luciferase was synthesized (Eurofins Genomics).

### mRNA expression analysis by RT-qPCR

1 µg of total RNA was prepared by column purification using the NucleoSpin® RNA Kit (Macherey-Nagel; #740955.250) and transcribed into cDNA using 0.5 µl RevertAid Reverse Transcriptase (Fisher Scientific #EP0441), 4 µl 5x reaction buffer, 0.5 µl Random Hexamer Primer, 0.5 mM dNTP mix (10 mM) in a total volume of 20 µl at 25°C for 10 min, 42°C for 1 h and 70°C for 10 min. 1 µl of the reaction mixture was used to amplify cDNA using Taqman® Gene Expression Assays (0.25 µl) (Applied Biosystems) primarily for *ACTB* (#Hs99999903_m1), *GUSB* (#Hs99999908_m1), *GAPDH* (#Hs02758991_g1), *IL8* (#Hs00174103_m1), *NFKBIA* (#Hs00153283_m1), *CXCL2 (#*Hs00236966_m1), *RELA* (#Hs01042019_g1) and TaqMan® Fast Universal PCR Master Mix (Applied Biosystems; #4352042). All PCRs were performed as duplicate reactions on an ABI7500 Fast real time PCR instrument. The cycle threshold value (ct) for each individual PCR product was calculated by the instrument’s software and Ct values obtained for inflammatory/target mRNAs were normalized by subtracting the Ct values obtained for *GUSB* or *ACTB or GAPDH*. The resulting ΔCt values were then used to calculate relative fold changes of mRNA expression according to the following equation: _2-((Δct *stim*.)-(Δct *unst.*)) or 2-((Δct siRNA target.)-(Δct *siRNA* Luciferase))._

### Targeted siRNA screen

For the siRNA screens, cDNA was synthesized in cell lysates and amplified using the TaqMan^®^ PreAmp Cells-to-Ct Kit^TM^ (Applied Biosystems; #4387299) and TaqMan^®^ Gene Expression Assays (Applied Biosystems) following an adapted miniaturized protocol. The kit enables to perform gene expression analysis directly from small numbers of cultured cells without RNA purification by an intermediate pre-amplification step between reverse transcription and qPCR. 3×10^3^ HeLa cells were seeded in 48-well plates and cultured overnight (24 h). Transfection occurred as described above with downscaled reagent volumes as follows: 12.5 µl Opti-MEM™, 1 µl Hi-PerFect Transfection Reagent and 4 µl of siRNA mixture (1 µM, finally 20 nM). 38 HCI and RELA were targeted by pools of 3-4 gene-specific siRNAs and compared to siLuciferase, Hi-PerFect only (HP), or untreated control samples. 48 h after transfection, half of the cells per plate were treated for 1 h with IL-1α (10 ng/ml). Cells were harvested by trypsinization and transferred to RNase-free reaction tubes on ice. Cells were washed twice with ice-cold PBS and lysed in 12.5 μl lysis solution (DNase I was diluted at 1:100). After vortexing, the lysates were incubated for 5 min at room temperature. The reaction was stopped by adding 1.25 μl stop solution. After repeated mixing, samples were incubated for 2 min at room temperature. The reverse transcription was directly conducted on the lysates by mixing 4.5 µl lysate (or nuclease-free water as a control) with a 5.5 µl RT mixture that was prepared as follows: 5 μl of 2× RT-buffer and 0.5 μl of 20× RT enzyme mix. The reaction tubes were incubated in a thermal cycler at 37°C for 60 min, then at 95°C for 5 min to inactivate the RT enzyme. In the following step, the cDNA was pre-amplified using gene-specific primers contained in TaqMan® Gene Expression Assays. The Assays of interest were diluted 1:100 in TE buffer. Therefore, two pools of Assays were prepared, with pool 1 for target genes 1-19 and pool 2 for target genes 20-38. Both pools were additionally supplemented with Assays for three prototypical NF-κB target genes *IL8* (#Hs00174103_m1), *NFKBIA* (#Hs00153283_m1) and *CXCL2* (#Hs00236966_m1), two housekeeping genes *GUSB* (#Hs99999908_m1) and *GAPDH* (#Hs02758991_g1) and the positive control *RELA* (#Hs01042019_g1). The pre-amplification PCR mixtures of pool 1 or pool 2 were prepared as follows: 2.5 μl pool 1/pool 2 were used in a 10 μl reaction volume with 5 μl 2×TaqMan PreAmp MasterMix and 2.5 μl cDNA. Samples with siRNA targets 1-19 were supplemented with the mixture of pool 1 and samples with siRNA targets 20-38 were supplemented with the mixture of pool 2. Controls were supplemented with each of the two pre-amplification mixtures. The pre-amplification occurred in a thermal cycler at 95°C for 10 min, following 15 cycles at 95°C for 15 sec/60°C for 4 min. Prior to real-time PCR, the pre-amplification products were diluted 1:5 with TE buffer. The expression of the indicated target genes was determined by real-time PCR using the TaqMan® Fast universal PCR master mix and 7500 Fast Real-Time PCR System from Applied Biosystems. Based on Ct values, mRNA levels were quantified and normalized against *GUSB*. The effects of knockdowns were calculated separately for basal and IL-1α-inducible conditions against the luciferase siRNA.

### Microarray transcriptomics

HeLa cells were transiently transfected for 48 h with 20 nM siRNA mixtures against *RELA*, *ZBTB5*, *S100A8*, *S100A9* (series 1) or *RELA*, *GLIS2*, *TFE3*, *TFEB* (series 2) and a siRNA against *luciferase* (siLuc) as control as described above. Half of the cells were treated with IL-1α (10 ng/ml) for 1 h at the end of the incubation, and Agilent microarray analyses were performed from isolated total RNA using the NucleoSpin^®^ RNA Kit (Macherey-Nagel; #740955.250). Per reaction 200 ng RNA was amplified and Cy3-labeled using the LIRAK kit (Agilent; #5190-2305) following the kit instructions. The Cy3-labeled aRNA was hybridized overnight to 8 x 60K 60-mer oligonucleotide spotted microarray slides (Agilent Technologies; # G4851C design ID 072363). Hybridization and subsequent washing and drying of the slides were performed following the Agilent hybridization protocol. The dried slides were scanned at 2 µm/pixel resolution using the InnoScan is900 (Innopsys, Carbonne, France). Image analysis was performed with Mapix 9.0.0 software, and calculated values for all spots were saved as GenePix results files. Stored data were evaluated using the R software and the limma package from BioConductor. Mean spot signals were background corrected with an offset of 1 using the NormExp procedure on the negative control spots. The logarithms of the background-corrected values were quantile-normalized. The normalized values were then averaged for replicate spots per array. From different probes addressing the same NCBI gene ID, the probe showing the maximum average signal intensity over the samples was used in subsequent analyses. Genes were ranked for differential expression using a moderated t-statistic. Pathway analyses were done using gene set tests on the ranks of the t-values. Pathway annotations were obtained from KEGG (Kanehisa et al., 2016). The genes assigned to these annotations, including the signal intensity values (E values) and the differential expression between samples (log_2_ fold change, LFC) with the associated significance (-Log_10_ p value) were listed in an Excel file and used for further filtering steps as mentioned in the figure legends.

### Cell lysis and immunoblotting

For whole cell extracts cells were lysed in Triton cell lysis buffer (10 mM Tris, pH 7.05, 30 mM NaPPi, 50 mM NaCl, 1% Triton X-100, 2 mM Na_3_VO_4_, 50 mM NaF, 20 mM ß-glycerophosphate and freshly added 0.5 mM PMSF, 2.5µg/ml leupeptin, 1.0 µg/ml pepstatin, 1 µM microcystin) and incubated for 15 min on ice. Lysates were cleared by centrifugation at 10,000 x g/4°C/15 min.

For preparation of nuclear and cytosolic extracts, cells were suspended and pelleted (800 x g/4°C/5 min) in buffer A (10 mM Hepes, pH 7.9, 10 mM KCl, 1.5 mM MgCl_2_, 0.3 mM Na_3_VO_4_, 20 mM β-glycerophosphate, freshly added 200 μM leupeptin, 10 μM E-64, 300 μM PMSF, 0.5 μg/ml pepstatin, 5 mM DTT and 1 µM microcystin). The pellet was resuspended in buffer A containing 0.1% NP-40 and incubated for 10 min on ice. After centrifugation at 10,000 x g for 5 min at 4°C, supernatants were taken as cytosolic extracts. Nuclear pellets were resuspended in buffer B (20 mM Hepes, pH 7.9, 420 mM NaCl, 1.5 mM MgCl_2_, 0.2 mM EDTA, 25% glycerol, 0.3 mM Na_3_VO_4_, 20 mM β-glycerophosphate, freshly added 200 μM leupeptin, 10 μM E-64, 300 μM PMSF, 0.5 μg/ml pepstatin, 5 mM DTT, and 1 µM microcystin). After 30 min on ice, nuclear extracts were cleared at 10,000 x g for 5 min at 4°C, and supernatants were collected.

For preparation of cytosolic, nuclear soluble and chromatin extracts, cells were washed, suspended and pelleted (500 x g / 4°C / 5 min) in PBS. Pellets were suspended in fractionated lysis buffer I (20 mM HEPES (pH 8.0), 10 mM KCl, 1 mM MgCl_2_, 0,1% (v/v) Triton X-100, 20% (v/v) glycerol, freshly added 50 mM NaF, 1 μM microcystin, 1 mM Na_3_VO_4_, 1 x Roche protease inhibitor cocktail) on ice for 10 min. After centrifugation for 1 min at 2300 x g at 4 °C, supernatants were taken as cytosolic extracts (C). The pellet was resuspended in fractionated lysis buffer II (20 mM HEPES, 2 mM EDTA, 400 mM NaCl, 0,1% (v/v) Triton X-100, 20% (v/v) glycerol, freshly added 50 mM NaF, 1 μM microcystin, 1 mM Na_3_VO_4_, 1 x Roche protease inhibitor cocktail), incubated on ice for 20 min, and briefly mixed twice during this time. Centrifugation for 5 min at 20,400 x g and 4 °C separated the nuclear soluble fraction (N1) in the supernatant. The remaining pellet was resuspended in fractionated lysis buffer III (20 mM Tris (pH 7.5), 2 mM EDTA, 150 mM NaCl, 1% (w/v) SDS, 1% (w/v) NP-40, freshly added 50 mM NaF, 1 μM microcystin, 1 mM Na_3_VO_4_, 1 x Roche protease inhibitor cocktail) and sonicated (6 cycles, high power for 30 s on and 30 s off at 4 °C, Bioruptor NextGen (Diagenode)). This was followed by incubation for 30 min on ice and centrifugation for 5 min at 20,400 xg and 4 °C. The supernatant contained the chromatin bound nuclear fraction (N2).

Protein concentrations of all cell extracts were determined by the Bradford method (Carl Roth; ROTI®Quant; #K929.3), and ∼20-50 µg protein per sample was supplemented with reducing gel-loading buffer 4xRotiLoad (Carl Roth; #K929.3). Immunoblotting was performed essentially as previously described (Hoffmann et al., 2005). Proteins were separated on SDS-PAGE and electrophoretically transferred to PVDF membranes (Roti-PVDF (0,45μm); Carl Roth; #T830.1). After blocking with 5% dried milk in Tris/HCl-buffered saline/0.05% Tween (TBST) for 1 h, membranes were incubated for 12-24 h with primary antibodies (diluted 1:500-1:10,000 in 5% milk or BSA in TBST), washed in TBST and incubated for 1-2 h with the peroxidase-coupled secondary antibody (HRP-coupled anti-rabbit IgG (Dako, #P0448), HRP-coupled anti-mouse IgG (Dako; #P0447). After washing in TBST, proteins were detected by using enhanced chemiluminescence (ECL) systems from Merck Millipore (Immobilon Western Chemiluminescent HRP Substrate; #WBKLS0500) or GE Healthcare (Amersham ECL Western Blotting Detection Reagent; #RPN2106). Images were acquired and quantified using the ChemiDoc TouchImaging System (BioRad) and the software ImageLab V_5.2.1 (Bio-Rad). For visualization of biotinylated proteins, membranes were blocked with 5% BSA in Tris / HCl-buffered saline / 0.05% Tween (TBST) for 24 h at 4 °C and afterwards membranes were incubated for 1 h with HRP-Streptavidin (PerkinElmer; #NEL750001EA; diluted 1:5000 in 5 % BSA in TBST).

### Immunofluorescence coupled to *in situ* proximity ligation assay (Immuno-PLA)

Immunofluorescence (IF) was coupled to *in situ* proximity ligation assay (Immuno-PLA). For PLA the Duolink® PLA reagents were used (Sigma-Aldrich; #DUO92007, #DUO92006, #DUO92002). 9000 cells per channel, of parental or p65-deficient cells, were seeded in ibiTreat μ-Slides VI 0.4 (Ibidi; #80606). On the next day, cells were washed twice with 150 µl PBS for 5 min and fixed with 100 µl of 4% paraformaldehyde (in PBS) for 10 min at room temperature. Afterwards, 100 µl 0.1 M Tris/HCl pH 7.4 were added and incubated for 10 min at room temperature. Permeabilization was performed by adding 100 µl of a 0.005% saponin/0.1% Triton X-100/PBS solution for 10 min at room temperature. Permeabilized cells were washed twice with 150 µl PBS for 5 min and incubated overnight (24 h) with 40% glycerol/ PBS at room temperature. On the next day, nuclei were permeabilized by three cycles of freeze-and-thaw. Ibidi-slides were therefore kept in liquid nitrogen for 1 min and thawed until glycerol cleared up. After nuclear permeabilization, cells were washed twice with PBS for 5 min and embedded in 100 µl blocking solution which was incubated in a humidity chamber for 30 min at 37°C. The blocking solution was discarded and cells were incubated with 50 µl of the appropriate primary antibody mixture in a humidity chamber for 1 h at 37°C. The primary antibody mixture contained PLA and IF antibodies (anti-NF-κB p65 (Santa Cruz Biotechnology; #sc-8008, ms and Bethyl Lab.; #A303-945A, gt), anti-TFE3 (Sigma-Aldrich; #HPA023881, rb), anti-TFEB (Cell Signaling; #4240, rb), anti-GLIS2 (Thermo Fisher Scientific; #PA5-40314, rb) and anti ZBTB5 (Sigma, #HPA021521, rb) was diluted in antibody diluent. Cells were washed three times with 150 µl buffer A for 5 min and then incubated with 50 µl of a secondary antibody mixture in a humidity chamber for 1 h at 37°C. The secondary antibody mixture contained PLA probes which were diluted 1:5 in antibody diluent but also the Dylight 488-coupled secondary anti ms IF antibody. From the time of incubation, all further steps were carried out in the dark. Cells were washed three times with 150 µl buffer A for 5 min and incubated with 50 µl ligase solution in a humidity chamber for 30 min at 37°C. The ligase was therefore diluted 1:60 in ligase solution (stock solution was diluted 1:5 in nuclease-free water). Cells were washed three times with 150 µl buffer A for 2 min and then incubated with 50 µl polymerase solution in a humidity chamber for 100 min at 37°C. The polymerase was therefore diluted 1:80 in amplification solution (stock solution was diluted 1:5 in nuclease-free water). Cells were first washed three times with 150 µl buffer B for 5 min followed by two HBSS washes for 1 min. Nuclear DNA was then stained with 1 µM Hoechst 33342 for 5 min at room temperature and washed twice with 150 µl HBSS for 5 min. Cells were finally embedded in 50 µl 30% glycerol/HBSS and stored in the dark until microscopic documentation. Fluorescence imaging of Immuno-PLA samples was carried out using the inverse fluorescence microscope THUNDER imager DMi8 (Leica Microsystems CMS GmbH; HC PL APO 20x/0.8 DRY objective, camera Leica-DFC9000GT-VSC13705) was used with filter cubes suited for Hoechst (excitation 391/32 and emission 435/30) and DyLight488 (excitation 506/21 and emission 539/24) with the Leica LASX software (version 3.7.4.23463). The Quantification of protein-protein interaction (PPI) complexes was performed using the Blobfinder software from the Centre for Image Analysis (Uppsala University, Sweden) and Olink Bioscience (Allalou and Wahlby, 2009). The software configuration was adjusted to HeLa cells with a nucleus size of 100 pixels^2^ and cytoplasm size of 100 pixels. For the blob size, the 3×3 default setting was applied whereas the blob threshold of 15 was determined by a test image.

### Chromatin immunoprecipitation (ChIP)

1.25-2.5×10^7^ KB cells were seeded in T175 cell culture flask per condition. At the next day they were starved for 24 h in HBSS or left untreated. After 23 h of starvation cells were stimulated with IL-1α (10 ng/ml) for 1 h or left unstimulated. Proteins bound to DNA were cross-linked *in vivo* with 1% formaldehyde added directly to the medium. After 10 min incubation at room temperature, 0.1 M glycine was added for 5 min to stop the cross-linking. Then, cells were collected by scraping and centrifugation at 1,610 x g (5 min, 4°C), washed in cold PBS containing 1 mM PMSF and centrifuged again. Cells were lysed for 10 min on ice in 3 ml ChIP lysis buffer (1% SDS, 10 mM EDTA, 50 mM Tris pH 8.1, 1 mM PMSF, 1,5x Roche protease inhibitor mix). The DNA was sheared by sonication (4x 7 cycles, 30s on / 30s off, power high; Bioruptor, Diagenode) at 4°C and lysates cleared by centrifugation at 16,100 x g at 4°C for 15 min. Supernatants were collected and stored in aliquots at –80°C. For determination of DNA concentration 20 μl of sheared lysate was diluted with 100 μl TE buffer including 10 μg/ml RNAse A. After 30 min at 37°C, 3.8 μl proteinase K (20 mg/ml) and 1% SDS was added and incubated for at least 2 h at 37°C followed by overnight incubation at 65°C for re-crosslinking. Samples were resuspended in two volumes of buffer NTB (Macherey-Nagel; #740595.150) and DNA was purified using the NucleoSpin^®^ Gel and PCR Clean-Up Kit (Macherey-Nagel; #740609.50) according to the manufacturer’s instructions. DNA was eluted with 50 μl 5 mM Tris pH 8.5 and concentration was determined by Nano Drop. For ChIP, the following antibody amounts were used: anti-NF-κB p65 (3 μg, Santa Cruz; #sc-372), anti-TFE3 (2 µg, Sigma-Aldrich; #HPA023881) and IgG (3 μg, Cell Signaling; #2729). Antibodies were added to precleared lysate volumes equivalent to 15 μg of chromatin. Then, 900 μl of ChIP dilution buffer (0.01% SDS, 1.1% Triton X-100, 1.2 mM EDTA, 167 mM NaCl, 16.7 mM Tris/HCl pH 8.1) were added and the samples were rotated at 4°C overnight. Thereafter, 30 μl of a protein A/G sepharose mixture (GE Healthcare; #17-0780-01 and #17-0618-01), pre-equilibrated in ChIP dilution buffer was added to the lysates and incubation continued for 2 h at 4°C. Beads were collected by centrifugation, washed once in ChIP low salt buffer (0.1% SDS, 1% Triton X-100, 2 mM EDTA, 20 mM Tris pH 8.1, 150 mM NaCl), once in ChIP high salt buffer (0.1% SDS, 1% Triton X-100, 2 mM EDTA, 20 mM Tris pH 8.1, 500 mM NaCl), once in ChIP LiCl buffer (0.25 M LiCl, 1% NP40, 1% desoxycholate, 1 mM EDTA, 10 mM Tris pH 8.1) and twice in ChIP TE buffer (10 mM Tris pH 8.1, 1 mM EDTA) for 5 min at 4°C end over end rotating. Beads were finally resuspended in 100 μl TE buffer including 10 μg/ml RNAse A. In parallel, 1/10 volume of the initial lysate (10% input samples) were treated accordingly. After 30 min at 37°C, 3.8 μl proteinase K (20 mg/ml) and 1% SDS were added and both, input and immunoprecipitates were incubated for at least 2 h at 37°C followed by overnight incubation at 65°C for re-crosslinking. Samples were resuspended in two volumes of buffer NTB and DNA was purified using the NucleoSpin^®^ Gel and PCR Clean-Up Kit. DNA was eluted with 50 μl 5 mM Tris pH 8.5 and stored at –20°C until further use. PCR products derived from ChIP-DNA were quantified by real-time PCR using the Fast ABI 7500 instrument (Applied Biosystems). The reaction mixture contained 2 µl of ChIP or input DNA (diluted 1:10 to represent 1% of input DNA), 0.25 µM of specific primers and 10 µl of Fast SYBR™ Green PCR Master Mix (Applied Biosystems; #4385612) in a total volume of 20 µl. PCR cycles were as follows: 95°C (20 sec), 40x (95°C (3 sec), 60°C (30 sec)). Melting curve analysis revealed a single PCR product. Calculation of enrichment by immunoprecipitation relative to the signals obtained for 1% input DNA was performed based on the equation % input = 2^-(Ct^ ^IP-Ct^ ^1%^ ^input)^. A list of oligonucleotides is provided in Supplementary Table 1.

### Motif analyses of ChIPseq data sets

NF-κB p65 ChIPseq peaks were compiled from four previously described data sets (Jurida et al., 2015) and were searched for enrichment of position-weight TF matrices using MEME-ChIP (https://meme-suite.org/) (Ma et al., 2014). Overrepresentation of TF motifs within ± 500 bp flanking the experimentally determined p65 / RELA ChIPseq peaks were calculated against the background of the whole human genome sequence (HG19) and is indicated by p value. The 1 kb windows were further searched for motifs of the TFs RELA, REL, TFE3, TFEB, GLIS2 and various ZBTB factors and predicted binding regions were annotated to genomic features to localize the next adjacent gene. The resulting matrices were filtered to assign motifs to the genes affected by siRNA knockdowns of TFs as indicated in the legends.

### Quantification and statistical analysis

Protein bands detected by Western blotting were quantified using Bio-Rad Image Lab, version 5.2.1 build 11. Statistics (t-tests, Mann-Whitney-Rank Sum Test, one-way ANOVA, correlations) were calculated using GraphPadPrism 9.5.1, Perseus 1.6.14.0 or Microsoft Excel 2016. PLA spots were quantified by Blobfinder (Allalou and Wahlby, 2009).

## Data availability

The proteomic data sets of this study have been submitted to the ProteomeXchange Consortium via the PRIDE partner repository (Perez-Riverol et al., 2022) with the dataset identifier PXD045888.

The microarray data sets of this study have been deposited in NCBI’s Gene Expression Omnibus (Edgar et al., 2002)and are accessible through GEO Series accession number GSE244637 (https://www.ncbi.nlm.nih.gov/geo/query/acc.cgi?acc=GSE244637).

For KB cells, RNA-seq, ChIP-seq and ATACseq data are available via our previous NCBI GEO submissions with the accession numbers GSE64224, GSE52470 and GSE134436 (https://www.ncbi.nlm.nih.gov/geo) (Jurida et al., 2015; Weiterer et al., 2020).

The remaining data generated in this study are provided in the Supplementary Information / Source Data sections. Source data are provided with this paper.

## Author contributions

L.L. and J.J. designed, performed and analysed TurboID, RNAi and the follow-up validation experiments and prepared graphs and tables, L.J. performed, analysed and visualized ChIP-qPCR experiments, L.L., J.J., L.J. and J.M-S. assembled the method section, C.M-B. helped with design and evaluation of PLAs, M.L.S. helped with p65-HA-mTb cloning, D.H. helped with cell selection experiments, A.W. processed LC-MS / MS raw data, U.L. performed LC-MS / MS analyses, M.B. re-analyzed ChIPseq data from L.J., J.W. performed microarray experiments and processed raw data, A.P. edited the first version of the manuscript, M.K. conceived the study, performed bioinformatics analyses, prepared figures and tables and wrote the initial draft, all authors contributed to the final version.

## Acknowledgments

This work was supported by the following grants from the Deutsche Forschungsgemeinschaft (DFG, German Research Foundation):

SFB1213/2 (B03 [to M.K., S.R., project 268555672]; TRR81/3 (A07 (to M.L.S.), B02 (to M.K.), project 109546710); KR1143/9-2 (KFO309, P3, project 284237345);

SFB1021/3 (C02 [to M.K.], Z03 [to M.K., U.L], project 197785619); and GRK2573 (RP5 [to M.K.], project 416910386).

Work in the laboratories of M.K. is also supported by the LOEWE program of the state of Hesse (Coropan, P4 to MK), the IMPRS program of the Max Planck Society and the Excellence Cluster CardioPulmonary Institute (EXC 2026: Cardio-Pulmonary Institute (CPI), project 390649896) and the DZL/UGMLC/ILH program.

---

**Supplementary Fig. 1.**
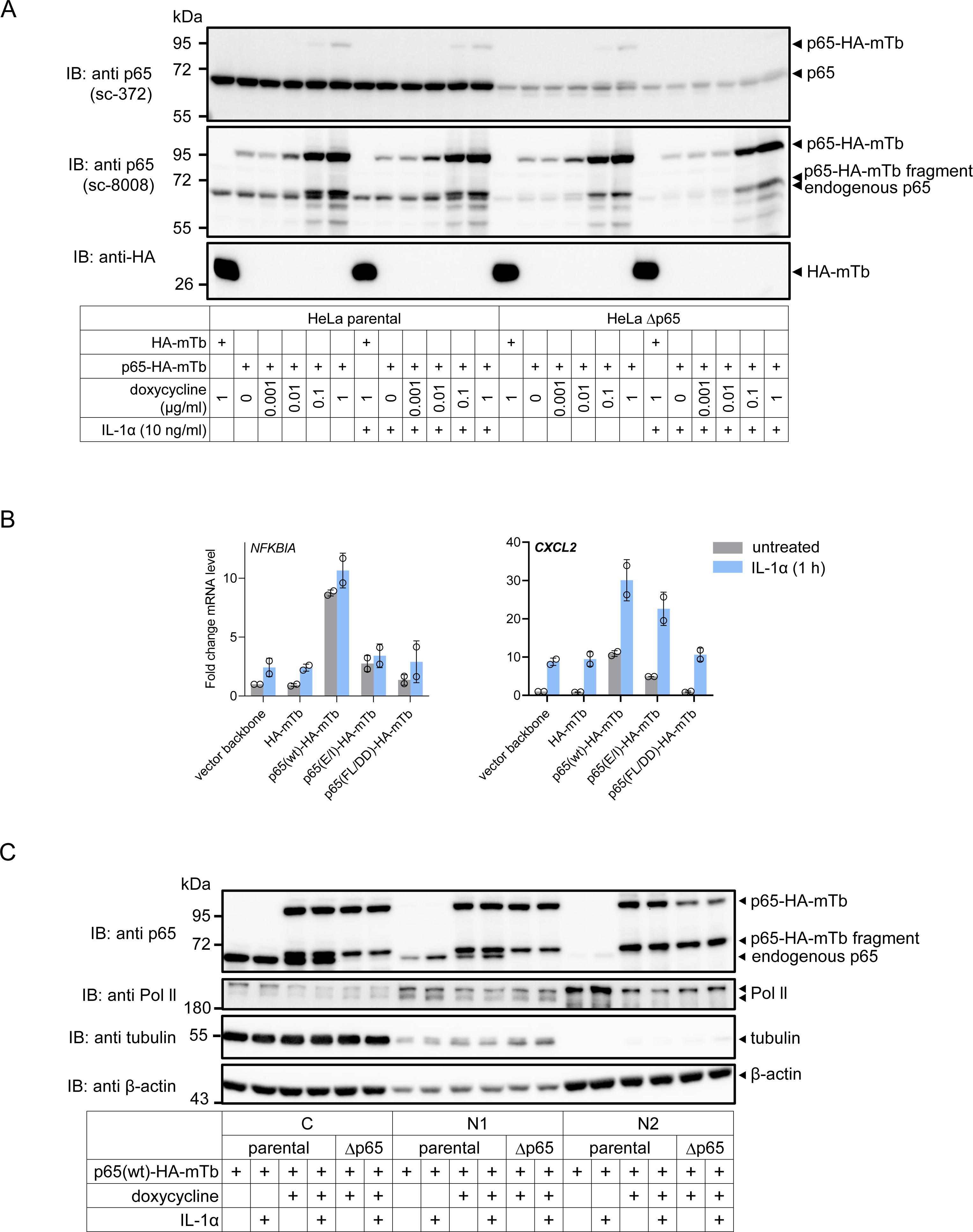
The p65-HA-miniTurbo fusion protein is inducibly expressed and functional. (A) Parental HeLa cells or pools of HeLa cells with CRISPR / Cas9-based suppression of endogenous p65 / RELA (Δp65) were transiently transfected with empty vector (EV) encoding HA-miniTurbo (HA-mTb) or with p65 / RELA wild type (wt) fused C-terminally to HA-mTb (p65(wt)-HA-mTb) as described in the legend of Fig. 1A. The expression of the constructs was induced with increasing concentrations of doxycycline for 17 h as indicated. At the end of the incubation, half of the cell cultures were treated with IL-1α (10 ng / ml) for 1 h. Cell extracts were analyzed by Western blotting for the expression of the p65-HA-mTb fusion protein or HA-mTb using polyclonal antibodies raised against the C-terminus of p65 / RELA (sc-372) or a monoclonal antibody raised against N-terminal amino acids 1-286 of p65 / RELA (sc-8008), or an anti HA antibody, respectively. Note that the fusion protein is better recognized with the N-terminal antibody preparations. (B) HeLa cells with CRISPR / Cas9-based suppression of endogenous p65 / RELA (Δp65) were transiently transfected with the indicated constructs and their expression was induced with doxycycline at 1 µg / ml for 17 h. On the next day, half of the cell cultures were treated with IL-1α (10 ng / ml) for 1 h. Total RNA was isolated and analyzed by RT-qPCR for expression of the indicated genes. Bar graphs show means ± s.d. from two biologically independent experiments. (C) Cells were transfected as in (A) and expression of the p65 / RELA fusion protein was induced 20 h later with doxycycline (10 ng / ml) for 4 h. In last period of this incubation, half of the cell cultures were treated with IL-1α (10 ng / ml) for 1 h. Cells were lysed and cytosolic (C), soluble nuclear (N1) and insoluble, chromatin nuclear fractions (N2) were analyzed by Western blotting for the expression and distribution of p65(wt)-HA-mTb. Antibodies against RNA polymerase II, tubulin and β-actin were used to control purity of fractions and equal loading.

**Supplementary Fig. 2.**
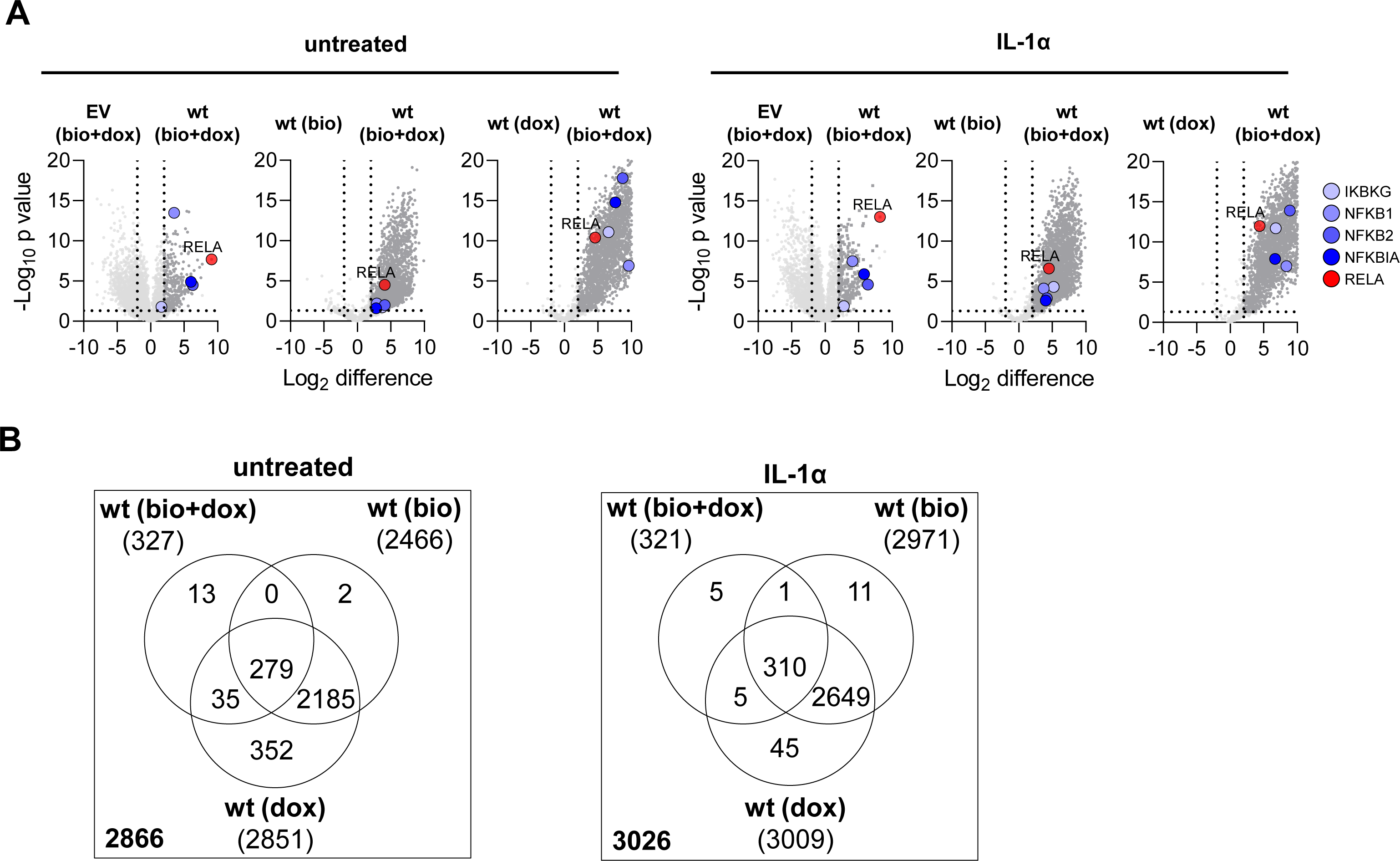
Identification of p65 / RELA high confidence interactors. (A) Biotinylated proteins from the experiments shown in Fig. 1C and from a second biological replicate were identified by mass spectrometry in the presence or absence of IL-1α treatment of cells. Volcano plots show the ratio distributions of Log_2_ transformed mean protein intensity values obtained with wild type p65 in the presence of doxycycline and biotin (wt) compared to the empty vector control (EV) or compared with conditions in which only biotin (wt(bio)) or doxycycline (wt(dox)) were added to the cell cultures, to determine false positive values in the absence of expression of fusion protein but facilitated biotinylation, or in the absence of biotinylation but induced expression of the fusion protein, respectively. X-axes show mean ratio value and Y-axes show p values from t-test results. Strong enrichment of the bait p65 / RELA proteins together with the core canonical NF-kB components is shown in red and blue colors, respectively (two biologically independent experiments and three technical replicates per sample). (B) Specific proteins binding to p65 / RELA wild type were defined by significant enrichment (LFC ≥ 2, -log_10_ p ≥ 1.3) compared to HA-miniTurbo only and to cells exposed to doxycycline or biotin only as shown in (A). Venn diagrams show the total numbers of specific p65 / RELA interactors and their overlaps before and after IL-1α-treatment. The intersecting 279 (without IL-1α) and 310 (with IL-1α) interactors were pooled, resulting in the set of 366 specific p65 / RELA interactors that was used for further downstream analyses. Numbers in the left lower corner of the boxes indicate the total number of detected interactors.

**Supplementary Fig. 3.**
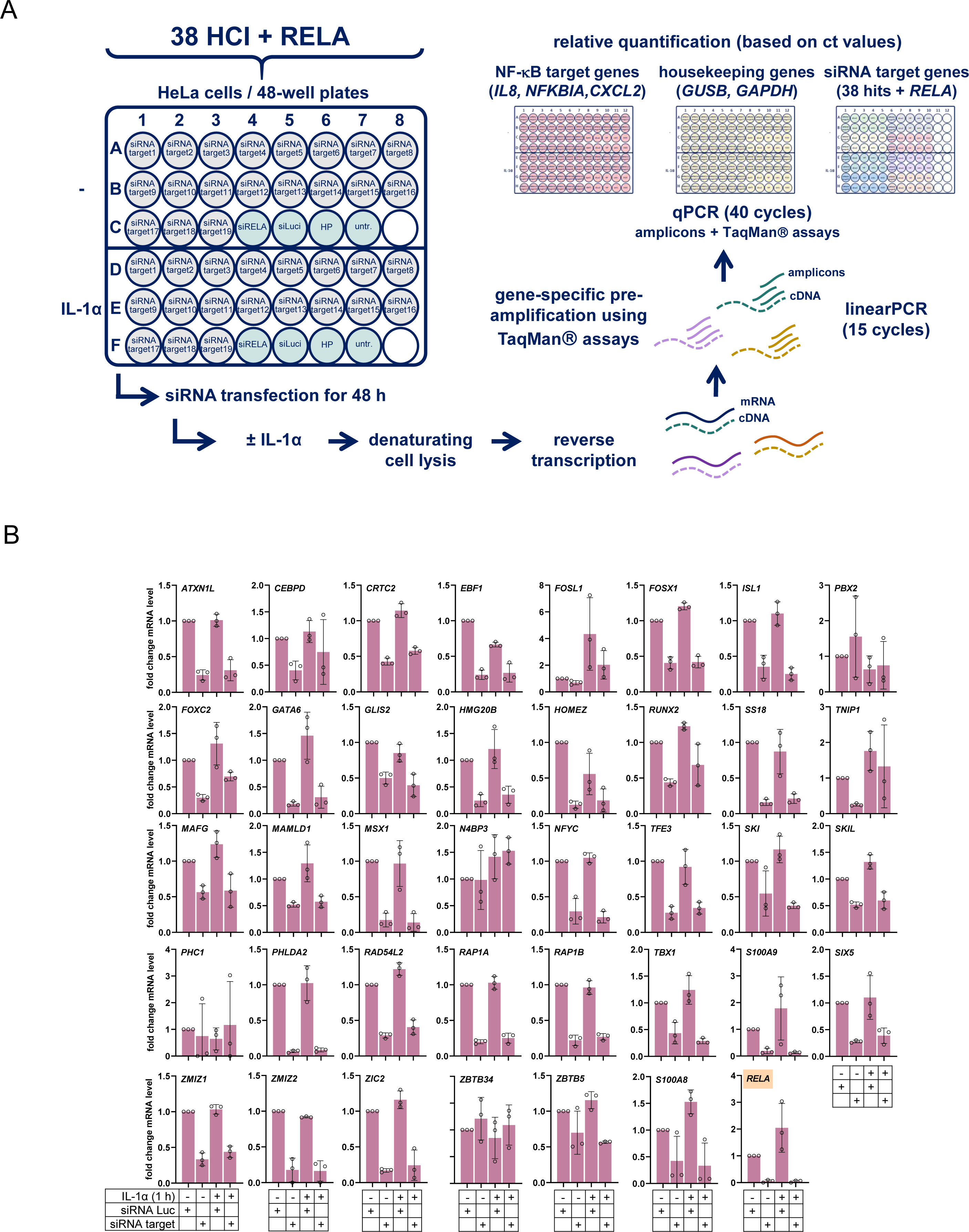
Targeted siRNA screen of 38 p65 / RELA high confidence interactors. (A) Scheme illustrating the arrangement of siRNAs and controls on individual cell culture plates and the performance of RT-qPCR measurements in cell extracts without prior RNA purification. A linear PCR amplification step was included to pre-amplify specific transcripts. (B) Confirmation of knockdown of 38 HCI and of RELA mRNAs by RT-qPCR as shown in (A). Bar graphs show mean changes ± s.d. relative to the luciferase siRNA controls (siLuci) from three biologically independent experiments.

**Supplementary Fig. 4.**
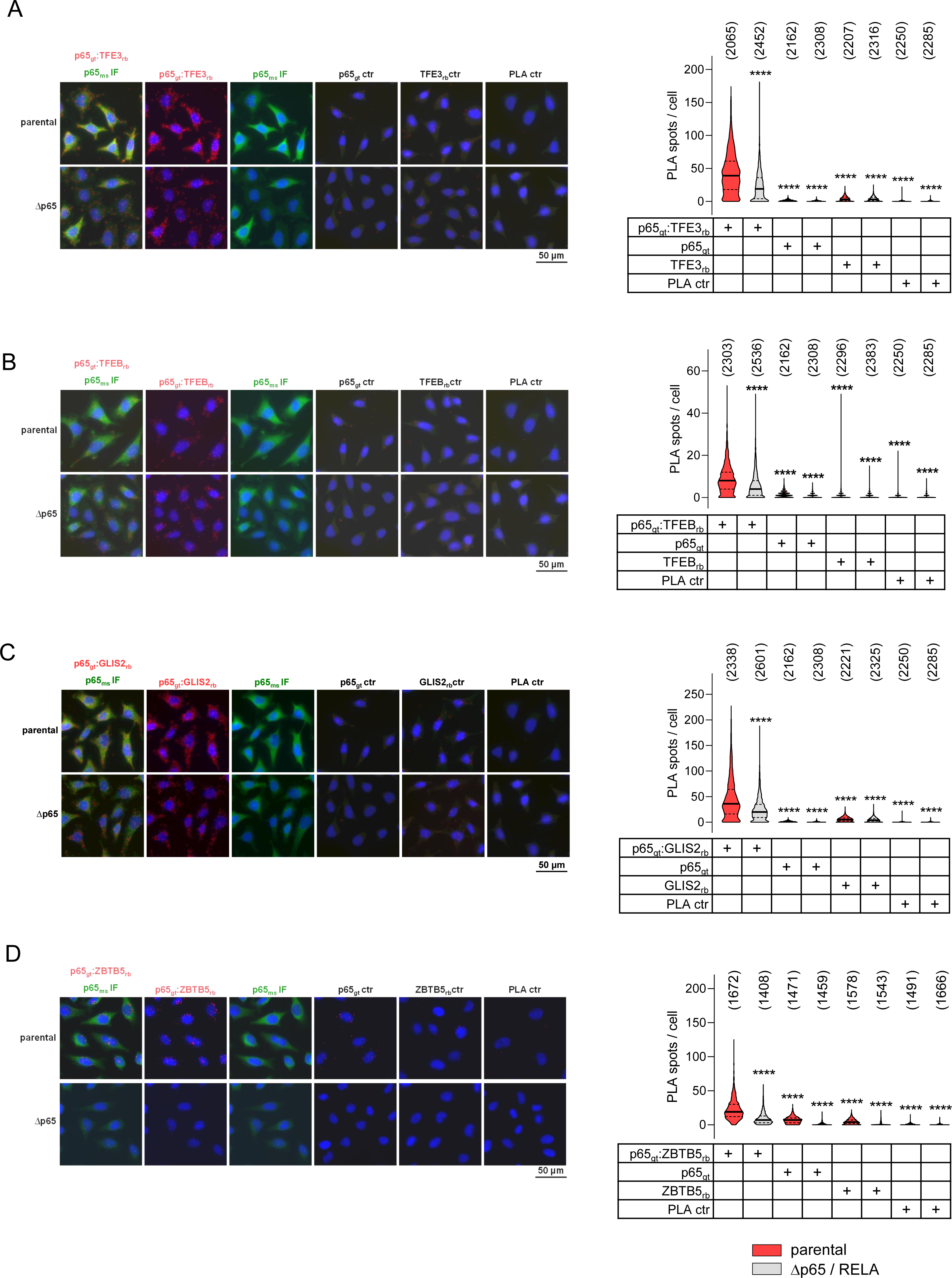
Proximity ligation assays confirm endogenous protein-protein interactions of p65 / RELA with interactors. Proximity-ligation assays coupled to immunofluorescence (IF) were performed with HeLa cells or Δp65 HeLa cells lacking endogenous p65 / RELA to demonstrate interactions of p65 / RELA with TFE3 (A), TFEB (B), GLIS2 (C) and ZBTB5 (D) using pairs of antibodies as indicated. PLA-spots are colored in red, while p65 IF is colored in green. Nuclear DNA is counterstained with Hoechst (blue signals). The images show representative fluorescence raw data and the violin plots on the right show quantification from the numbers of cells indicated in brackets. Samples omitting one of the two antibodies or both primary antibodies (ctr) served as negative controls. Solid lines indicate medians and dashed lines indicate 1^st^ and 3^rd^ quartiles. Asterisks indicate results from Kruskal-Wallis tests compared to the parental control (****p ≤ 0.0001). obtained by one-way ANOVA.

**Supplementary Fig. 5.**
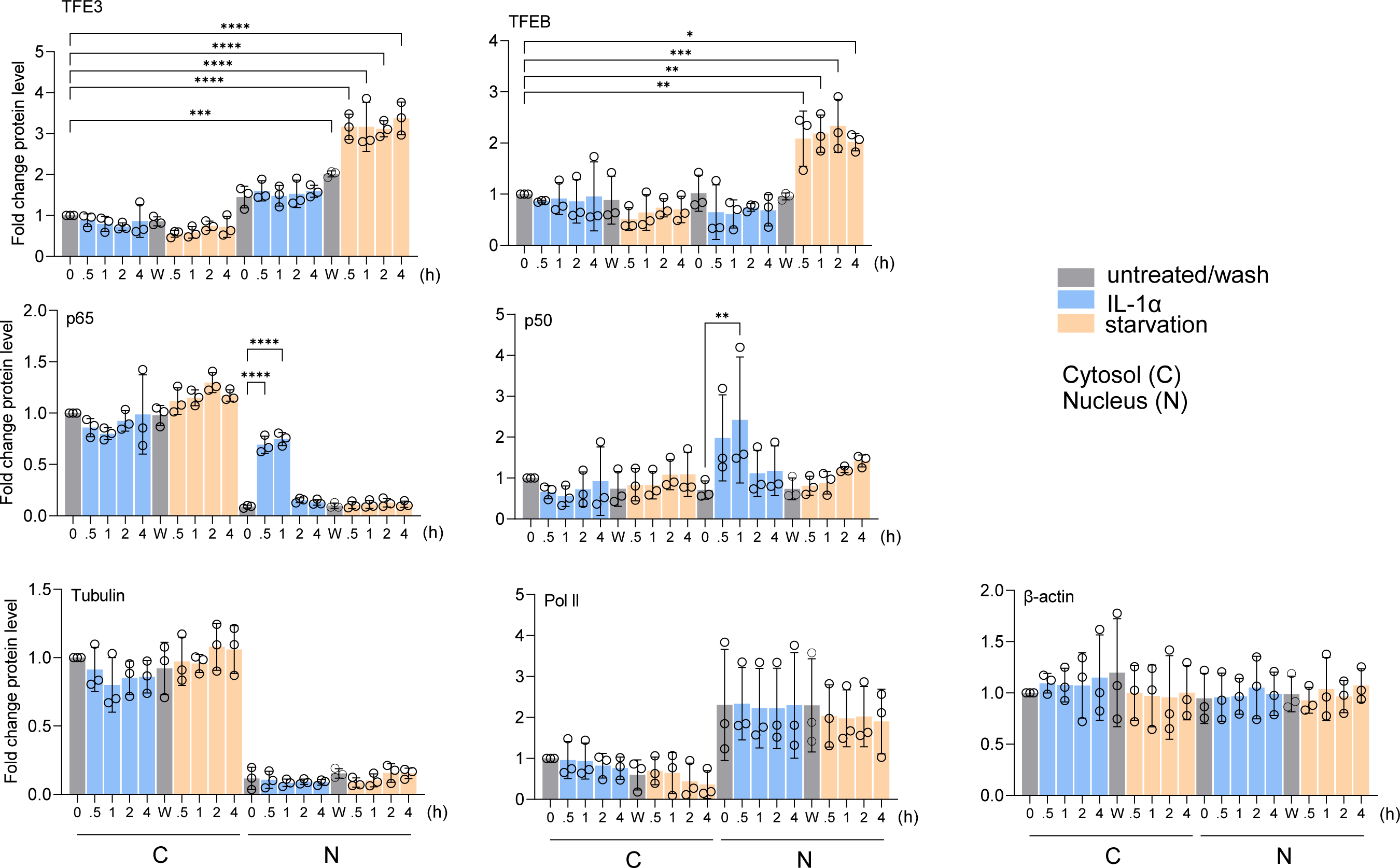
Quantification of subcellular distributions of p65 / RELA, TFE3 and TFEB. Cells were treated and cell extracts were analyzed by Western blotting as described in the legend of Fig. 4B. Bar graphs show mean changes ± s.d. relative to untreated controls from three independent experiments. Asterisks indicate p values (*p ≤ 0.05, **p ≤ 0.01, ***p ≤ 0.001, ****p ≤ 0.0001) obtained by one-way ANOVA. C = cytosol; N = nucleus.

**Supplementary Fig. 6.**
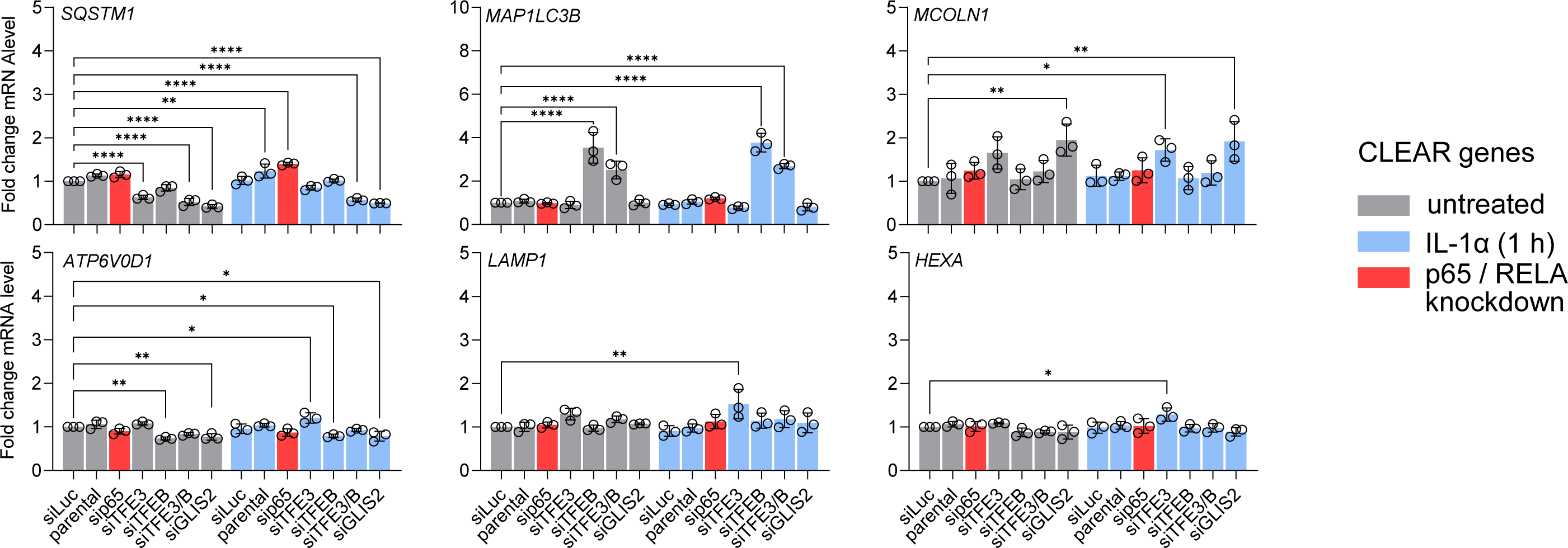
CLEAR gene expression does not depend on p65 / RELA. Total RNA isolated from cells treated as in Fig. 4C was analyzed for mRNA expression of the indicated CLEAR target genes by RT-qPCR. Data show mean values relative to cells transfected with luciferase siRNA ± s.d. from three biologically independent experiments. DK indicates double knockdown of TFE3 and TFEB. Asterisks indicate p values (*p ≤ 0.05, **p ≤ 0.01, ***p ≤ 0.001, ****p ≤ 0.0001) obtained by one-way ANOVA.

**Supplementary Fig. 7.**
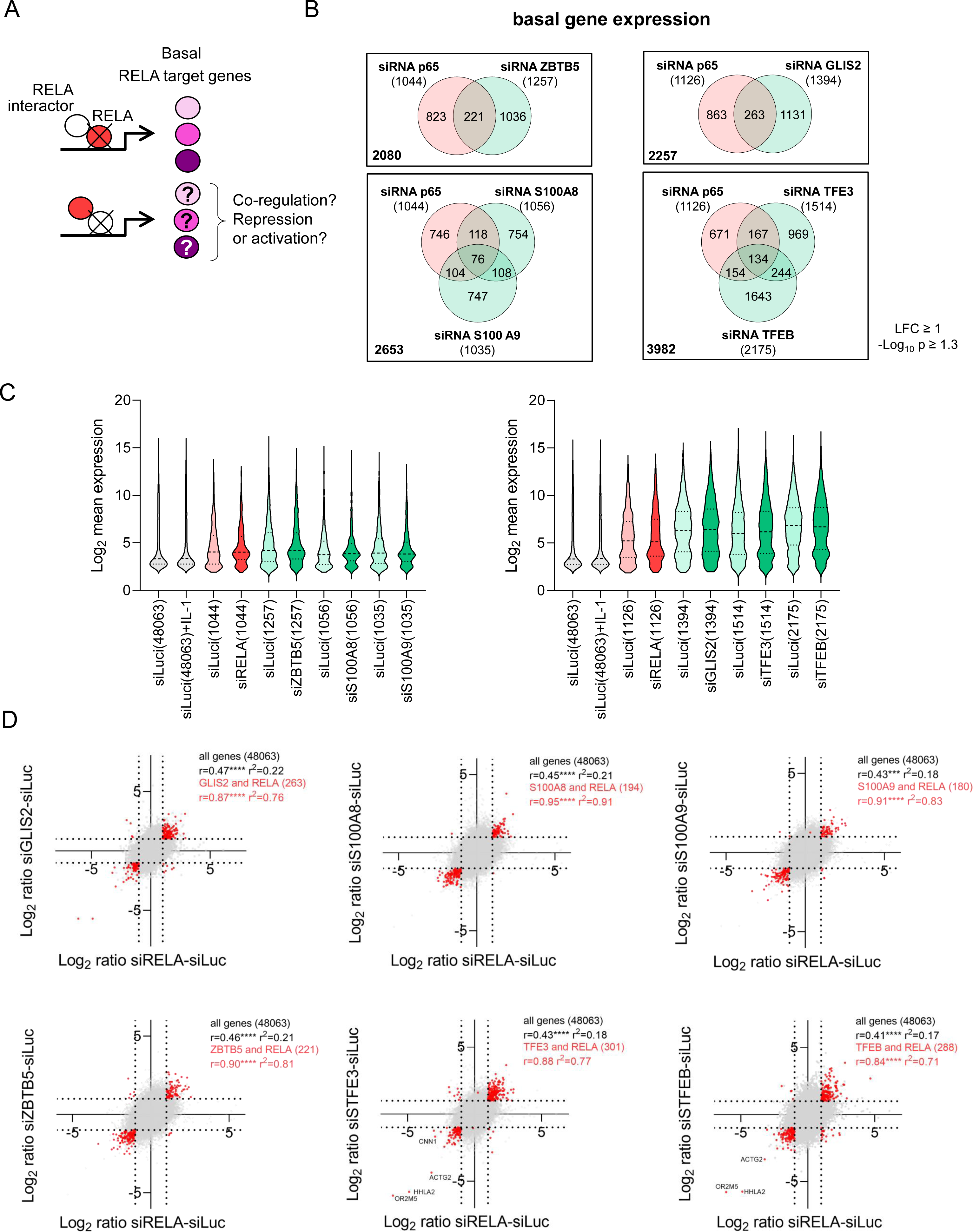
ZBTB5, GLIS2, S100A8 / S100A9 and TFE3 / TFEB co-regulate constitutively expressed subsets of p65 / RELA target genes. (A) Schematic illustrating the strategy to analyze the influences of novel p65 / RELA interactors on basal p65 / RELA target genes by combining siRNA-mediated knockdown with transcriptome analysis. (B) HeLa cells were transiently transfected for 48 hours with 20 nM siRNA mixtures against RELA, ZBTB5, S100A8, S100A9 (series 1) or RELA, GLIS2, TFE3, TFEB (series 2) and an siRNA against luciferase (siLuc) as control. Half of the cells were treated with IL-1α (10 ng/ml) for 1 hour at the end of incubation, and Agilent microarray analyses were performed from total RNA. Normalized data were used to identify DEGs based on an LFC ≥ 1 with a -log_10_ p value ≥ 1.3. Venn diagrams show the overlap of all DEGs that were affected at least twofold by siRNA knockdown in untreated, basal conditions, with the ratio of siLuc to individual knockdown determined in each case. Red colors mark genes jointly regulated by knockdown of RELA and one of its interactors (two biologically independent experiments). (C) Violin plots show the distribution, medians, and interquartile ranges of normalized expression levels for all constitutively expressed genes and the corresponding changes in the gene subsets defined in Supplementary Fig. 7B that were affected by siRNA knockdown. The number of these genes is indicated in parentheses. (D) Superimposed pairwise correlation analyses of the mean ratio changes of all genes (gray), and gene sets significantly up- or down-regulated by siRNA knockdown (red). Ratio values from RELA knockdown conditions were compared with the knockdown of a RELA interactor in each case. Genes that are jointly regulated by knockdown of RELA and one of its interactors correspond to the Venn diagrams of (B) and are marked in red. Coefficients of correlation (Pearson’s r), corresponding p values and coefficients of determination (r^2^) rare indicated for all comparisons. The complete set of data is provided in Supplementary Table 3.

**Supplementary Fig. 8.**
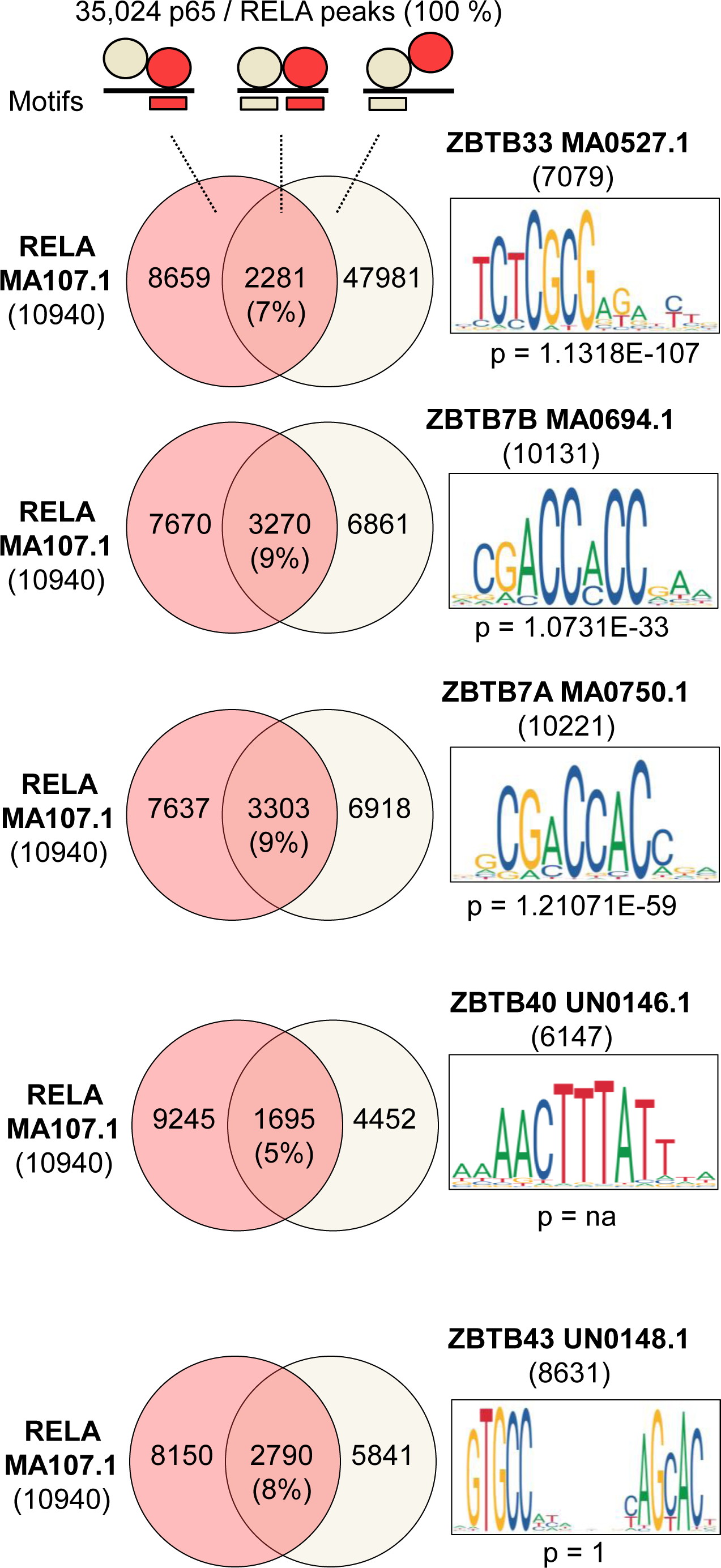
Motif analyses of ZBTB factors. Venn diagrams indicating the overlap of RELA motifs with motifs of ZBTB factors that were found by miniTurboID to interact with RELA, in chromosomal regions assigned to p65 / RELA ChIPseq peaks. P values indicated significant enrichment compared to the whole genome. Inserts show motif compositions.

**Supplementary Table 1. Source_data_of_LC-MS_MS_experiments.**

**Supplementary Table 2. Source_data_of_targeted_siRNA_screen.**

**Supplementary Table 3. Source_data_of_microarray_experiments.**

**Supplementary Table 4. Source_data_motif_analyses_under p65_RELA_ChIPseq_peaks.**

**Supplementary Table 5. NF-kB_interactors_from_published_large_scale_screens.**

